# KDM2B Silencing Elicits a Paracrine Mechanism Which Destabilizes SLUG, Promoting Differentiation of Basal-Like Breast Cancer Cells

**DOI:** 10.1101/2020.05.21.109819

**Authors:** Elia Aguado-Fraile, Burak Soysal, Vollter Anastas, Zeynep Tugce Soysal, Oksana Serebrennikova, Maria D. Paraskevopoulou, Evangelia Chavdoula, Philip N. Tsichlis

## Abstract

KDM2B is a JmjC domain H3K36me2/H3K36me1 demethylase, which immortalizes cells in culture and contributes to the biology of both embryonic and adult stem cells, including cancer stem cells. Here we show that the silencing of KDM2B activates a tyrosine kinase receptor-dependent paracrine mechanism, which results in the downregulation of *SNAI2* (SLUG), *SNAI1* (SNAIL) and SOX9, which also contribute to the biology of stem and progenitor cells. The downregulation of these molecules is posttranscriptional and in the case of *SNAI2*-encoded SLUG, it is due to calpain-dependent proteolytic degradation. SLUG abundance in normally growing cells is under the homeostatic control of GSK3, which phosphorylates SLUG and tags it for proteasomal degradation. The paracrine mechanism activated by KDM2B depletion, activates FGFR1 and EGFR family members, and blocks the homeostatic SLUG degradation by inactivating GSK3. This, however, sensitizes SLUG to classical calpains, which are also activated in KDM2B-depleted cells via Ca2+ influx and calpastatin downregulation. The switch in SLUG degradation pathways, results in the rapid degradation of SLUG and the differentiation of breast cancer stem cells, revealing an unexpected mechanism of stem cell regulation by a lysine demethylase.

**HIGHLIGHTS:**

- GSK3 phosphorylates SLUG and promotes its homeostatic proteasomal degradation
- KDM2B depletion results in GSK3 inactivation, Ca2+ upregulation and calpain activation.
- Downregulation of SLUG phosphorylation by GSK3, sensitizes SLUG to calpain activation.
- GSK3 inactivation and Ca2+ upregulation are due to an RTK-dependent paracrine mechanism

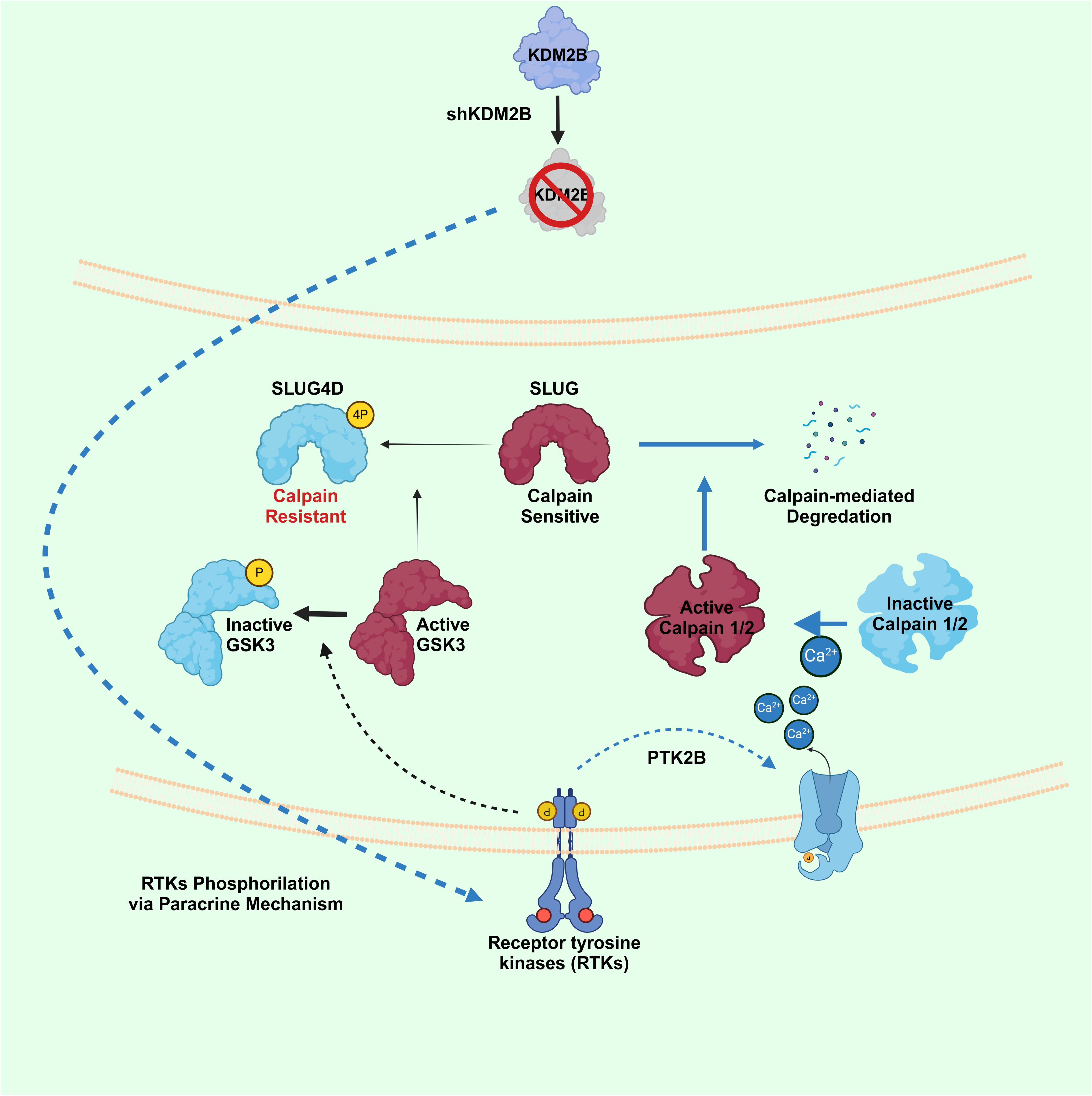

## INTRODUCTION

Basal-like breast cancer comprises 15-20% of all breast cancers and is more prevalent in younger women ^1^. These tumors do not express estrogen and progesterone receptors, and they are negative for epidermal growth factor receptor 2 amplification ^2^. Although the basal-like breast cancer subtype is heterogeneous ^3^, the majority of tumors in this subtype are characterized by an aggressive clinical course, early relapse after treatment and poor overall survival ^1,4^. The poor prognosis is partially due to the lack of effective targeted therapies for these tumors ^4^. The goal of the work presented in this report is to improve our understanding of the biology of this disease, which may lead to the development of novel and more effective therapeutic strategies. Its focus is on KDM2B, an enzyme which tends to be expressed at high levels in basal-like breast cancer, and whose expression is higher in a subset of tumors with an unfavorable metabolic phenotype ^5^.

*KDM2B* (also known as *NDY1, FBXL10, JHDM1B* or *Fbl10*), encodes a Jumanji C (JmjC) domain-containing lysine demethylase, which targets histone H3K36me2/me1 and perhaps histone H3K4me3 ^6–9^ and histone H3K79 me3/me2 ^10^. In addition to the JmjC domain, which is responsible for its demethylase activity ^7^, KDM2B contains CXXC and PHD zinc finger domains, an F-box and a leucine-rich repeat ^7,8^, and functions either as a transcriptional repressor, or a transcriptional activator ^5,11^. Transcriptional repression relies on the integration of multiple epigenetic signals, which is due to the KDM2B-dependent coupling of H3K36me2/me1 demethylation with H3K27 trimethylation and H2AK119 ubiquitination ^9,12–14^. Transcriptional activation on the other hand, depends on the noncanonical PRC1.1 complex, which is targeted to DNA by KDM2B, one of its components, and on MYC and ATF4, which bind the promoters of transcriptionally active genes in concert with KDM2B ^5,11^. These activities of KDM2B appear to depend on its demethylase activity. However, a variant of KDM2B lacking the JmjC demethylase domain also contributes to the function of KDM2B by inhibiting the methylation of a subset of CpG islands associated with bivalent developmental genes ^15,16^.

KDM2B functions as an oncogene in several types of tumors. Following the original discovery of its oncogenic potential in Moloney murine leukemia virus (MoMuLV)-induced rodent T cell lymphomas, where it was found to be activated by provirus integration ^7,9^, it was shown to function as an oncogene in human lymphoid ^17^ and myeloid malignancies ^18^ as well as in bladder cancer ^13,19,20^ and pancreatic cancer ^21^, basal-like breast cancer ^17,22^, gliomas ^17,23^ and prostate cancer ^20,24^. The oncogenic potential of KDM2B is mediated by multiple mechanisms. Our early studies revealed that, when overexpressed in normal mouse embryo fibroblasts, KDM2B stimulates cell proliferation and blocks replicative senescence by promoting the phosphorylation of Rb and by blocking the cell cycle inhibitory effects of p21^CIP1^ ^7,20^. Subsequently, we and others showed that the ability of KDM2B to stimulate the cell cycle and to block senescence may also be due to the repression of p16^INK4A^ and p15^INK4B^, which it achieves by acting in concert with EZH2 ^9,20,25^. Our studies addressing the effects of the KDM2B knockdown in a set of ten cancer cell lines, showed that in addition to the inhibition of the cell cycle, which was common to all, the knockdown of KDM2B also induced apoptosis in some ^20,22^.

Our early studies addressing the role of KDM2B in cellular metabolism revealed that KDM2B promotes the expression of a set of antioxidant genes, including aminoadipate-semialdehyde synthase (Aass), NAD(P)H quinone dehydrogenase 1 (Nqo1), peroxiredoxin 4 (Prdx4) and serpin family B member 1b (Serpinb1b), and protects cells from oxidative stress ^26^. Our more recent studies revealed that KDM2B MYC and ATF4 regulate in concert the expression of multiple metabolic enzymes, including the enzymes that control the interconnected SGOC, glutamate and GSH pathways ^5^. Importantly, the same transcriptional network controls the expression of a host of ribosomal protein and ribosome biogenesis factors and regulates the assembly and maturation of ribosomal subunits in the nucleolus. As a result, it controls the rate of assembly of new ribosomes, ribosome abundance and mRNA translation ^11^

KDM2B plays an important role in the self-renewal and differentiation of both normal and cancer stem cells. First, it’s expression tends to be higher in embryonic and adult stem cells than in other somatic cells. Its overexpression in somatic cells promotes reprogramming to induced pluripotent stem cells (iPSCs) ^20,27,28^ and its overexpression in fibroblasts contributes to reprogramming into hepatocyte-like cells ^20,29^. KDM2B also promotes the self-renewal of hematopoietic and chondrogenic stem cells ^17,20,30–32^ and the commitment of hematopoietic cells toward the lymphoid cell lineage ^17,20^. Our studies focusing on cancer stem cells, revealed that KDM2B is induced by FGF-2, and that it functions in concert with the histone H3K27 methyltransferase EZH2 to repress the expression of a set of microRNAs, which target multiple components of polycomb repressive complex 1 (PRC1) (BMI1 and RING1B), noncanonical polycomb repressive complex 1.1 (ncPRC1.1) (RING1B) and polycomb repressive complex 2 (PRC2) (EZH2 and SUZ12). The repression of these microRNAs results in the upregulation of their polycomb targets, promoting the self-renewal of cancer stem cells. The repression of the microRNAs that target the polycomb complexes depends on the demethylase activity of KDM2B, as evidenced by the fact that overexpression of a catalytically inactive KDM2B does not promote the recruitment of EZH2 to the miR-101 promoter and the repression of miR-101. However, since both KDM2B and EZH2 are required for the microRNA repressive function of KDM2B, EZH2 alone cannot rescue the PRC1 and PRC2 downregulation phenotype elicited by the knockdown of KDM2B ^13,20,22^.

The biology of mammary stem cells also depends on SLUG (*SNAI2*), SNAIL (*SNAI1*) and SOX9. Earlier studies had shown that SLUG is expressed in the basal myoepithelial layer of the adult mammary gland and that cells expressing SLUG possess stem cell properties ^20,33,34^. Additionally, transient overexpression of SLUG in luminal progenitor cells, sufficed to reprogram them into mammary stem cells (MaSCs) ^20,33^. Finally, differentiation of basal/myoepithelial cells into luminal cells was associated with the downregulation of SLUG ^20,33,35^. Other studies revealed that ectopic expression of SNAIL, a transcription factor closely related to SLUG, also contributes to the reprogramming of luminal cells into basal/myoepithelial cells ^20,33^. Given that KDM2B is also involved in the regulation of stem cell self-renewal and differentiation, these findings suggested a functional relationship between SLUG, SNAIL and KDM2B. The role of SOX9 in the regulation of mammary cell differentiation was suggested by in vivo studies showing that SOX9 is expressed in a subpopulation of cells in the basal/myoepithelial layer of the mammary ducts and that some of these cells express both SLUG and SOX9 ^20,33^. In addition, SOX9 complemented the ability of SLUG to reprogram fully differentiated luminal cells into cells with stem cell properties ^20,33^. Thus, whereas SLUG alone fails to reprogram fully differentiated mammary epithelia, the combined expression of SLUG and SOX9 in these cells gives rise to mammary stem cells which assemble to form solid organoids in Matrigel cultures and to produce a complete mammary ductal tree in the murine mammary gland reconstitution assay ^33^. Based on these considerations, we addressed the effects of the knockdown of KDM2B on the expression of SLUG, SNAIL and SOX9, and we observed that all three proteins are downregulated post transcriptionally by shKDM2B in human mammary gland-derived cell lines.

In this report we focus on SLUG, one of these factors, and we show that the knockdown of KDM2B promotes its degradation via a paracrine mechanism, which depends on FGFR1 and EGFR family members and activates calpains 1 and 2, by promoting intracellular Ca2+ upregulation and calpastatin downregulation. Importantly, the sensitivity of SLUG to calpain-mediated proteolytic degradation was enhanced by a block in GSK3-mediated SLUG phosphorylation, which was also induced by the shKDM2B-activated paracrine mechanism. The homeostatic regulation of SLUG in normally growing cells is under the control of GSK3, which phosphorylates SLUG and tags it for proteasomal degradation ^36^. The shKDM2B-induced paracrine mechanism, therefore, blocks the slow homeostatic mechanism of SLUG degradation, and switches to a rapid, calpain-dependent degradation mechanism, which results in the robust downregulation of SLUG, promoting mesenchymal to epithelial transition (MET) and the differentiation of mammary epithelial cell lines.

## RESULTS

### KDM2B regulates the expression of SLUG (*SNAI2*), SNAIL (*SNAI1*) and *SOX9* post transcriptionally

To determine whether KDM2B regulates the expression of SLUG, SNAIL and SOX9, we knocked down KDM2B in six immortalized or transformed, human mammary gland-derived cell lines and we examined the effects of the knockdown on the expression of these molecules, at both the RNA and the protein levels. This revealed a dramatic shKDM2B-induced downregulation of SLUG and SNAIL in all the cell lines and of SOX9 in some. Given that the RNA levels of these genes were upregulated rather than downregulated, we conclude that their downregulation was posttranscriptional (Fig 1A, 1B and S1A). In the following sections of this report, we will present mechanistic studies on the KDM2B-depndent posttranscriptional regulation of SLUG, which appears to be the critical regulator of the biology of mammary stem and progenitor cells ^20,33^.

**Figure 1:**
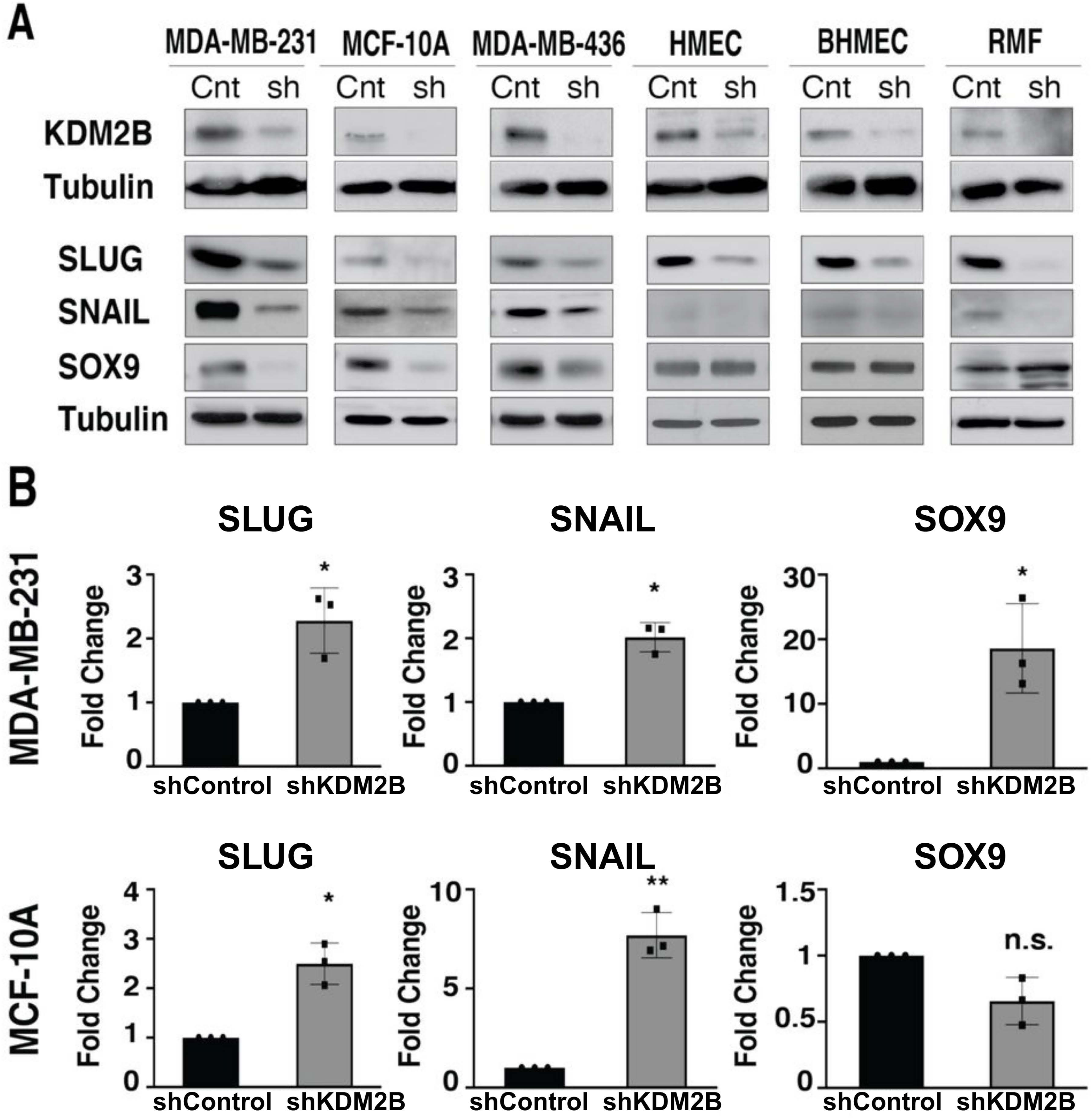
KDM2B regulates the expression of SLUG, SNAIL and SOX9 postranscriptionally: ***A.*** Immunoblotting of cell lysates of a panel of human mammary gland-derived cell lines transduced with shControl (Control) or shKDM2B (sh) lentiviral constructs, shows that the knockdown of KDM2B results in the downregulation of SLUG, SNAIL and SOX9. **B.** Quantitative RT-PCR failed to detect downregulation of SLUG, SNAIL and SOX9 in shKDM2B-transduced MDA-MB-231 and MCF10A cells. qRT-PCR of RNA from the remaining cell lines (MDA-MB-436, HMEC, BHMEC and RMF) gave similar results (Fig S1A)

### The downregulation of SLUG in KDM2B knockdown cells is not the result of mRNA translational repression

We recently showed that KDM2B regulates ribosomal biogenesis and translation and that the effect of the knockdown of KDM2B on the efficiency of translation varies between mRNA transcripts (68). This suggested that the shKDM2B-induced posttranscriptional downregulation of SLUG could be the result of transcript specific translational repression. To address this question, we searched our paired RIBO-Seq/RNA-Seq data in control and shKDM2B-transduced MDA-MB-231 cells ^11^ and we observed that that the translational efficiency of the SLUG mRNA was not affected by the knockdown of KDM2B (Fig S1C). To validate the results of this analysis, control and shKDM2B-transduced MDA-MB-231 cells were starved of methionine for 1 hour, and following this, they were cultured in methionine-free media supplemented with the methionine analog L-Homopropargylglycine (L-HPG). L-HPG contains an alkyne group, and its incorporation into newly synthesized proteins can be monitored with biotin azide, using “click chemistry”. Employment of this detection method, allowed us to monitor the abundance of SLUG in western blots of cell lysates, harvested at sequential time points from the start of the L-HPG pulse. Normalization of the abundance of SLUG, for the abundance of total protein in the lysates, confirmed that the rate of SLUG accumulation over time was not selectively affected by the knockdown of KDM2B (Fig S1D),

### KDM2B regulates the stability of SLUG

Based on the preceding data we hypothesized that the knockdown of KDM2B compromises the stability of SLUG. To address this hypothesis, we treated control and shKDM2B-transduced MDA-MB-231 and MCF10A cells with the protein synthesis inhibitor Cycloheximide (CHX) (30 μg/ml) and we monitored the protein levels of SLUG over time by western blotting. The results revealed that SLUG was indeed less stable in shKDM2B-transduced cells (Fig 2A and S1B).

**Figure 2:**
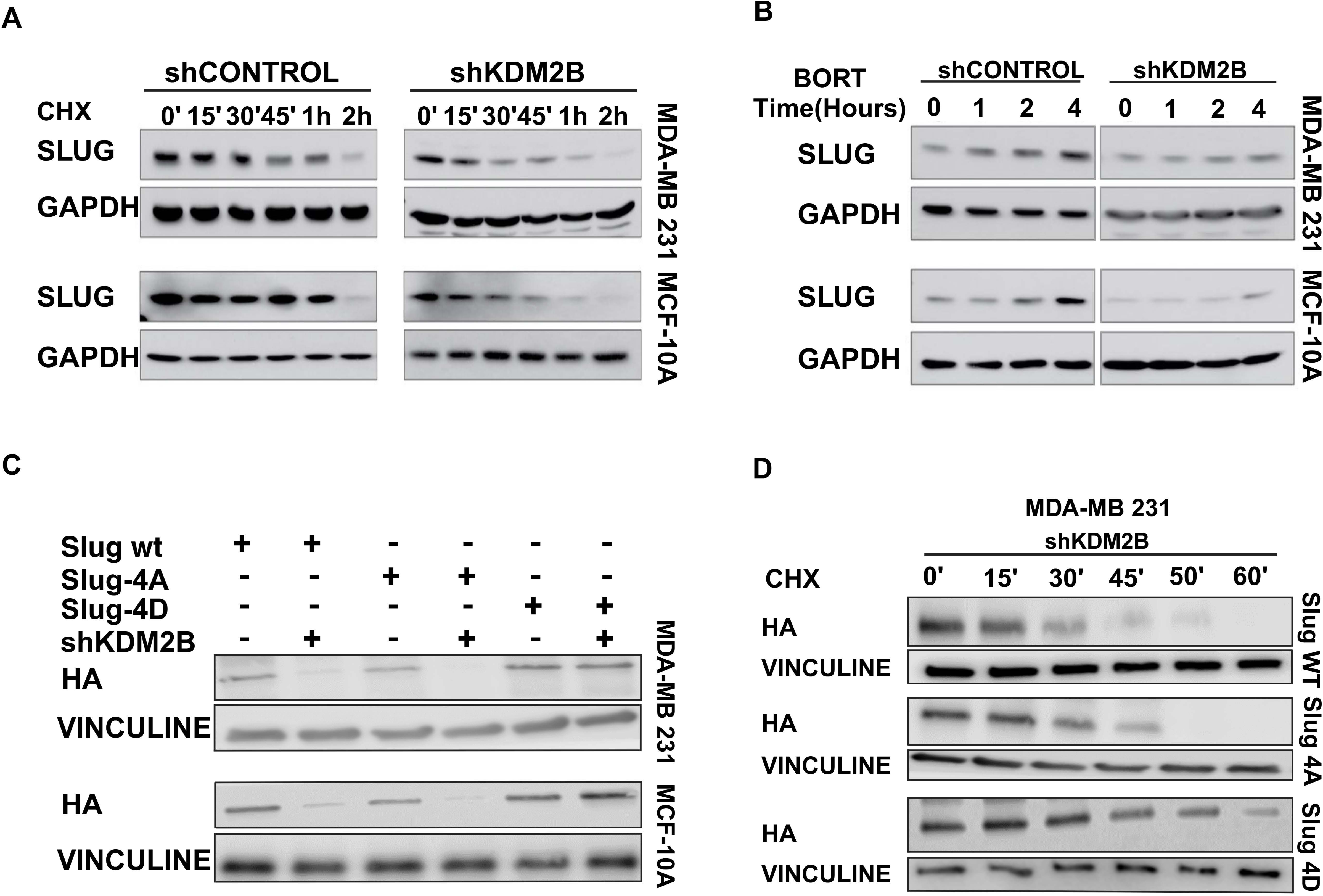
shKDM2B destabilizes SLUG by a proteasome-independent mechanism: **A.** shControl (Left panel) and shKDM2B-transduced (Right panel) MDA-MB-231 and MCF-10A cells were treated with Cycloheximide (CHX) (30 μg/ml) and they were harvested at sequential time points, as indicated. Probing western blots of the harvested cell lysates with anti-SLUG or anti-GAPDH (loading control) antibodies, showed that the degradation of SLUG was more rapid in the shKDM2B cells. **B.** shControl and shKDM2B-transduced MDA-MB-231 and MCF-10A cells were treated with the proteasome inhibitor Bortezomib (BORT) (0.5 μM) and they were harvested at sequential time points, as indicated. Probing western blots of the harvested cell lysates with anti-SLUG or anti-GAPDH (loading control) antibodies, showed a more robust upregulation in the shControl than in the ShKDM2B-transduced cells, suggesting that the destabilization of SLUG by shKDM2B is proteasome independent. **C.** shControl and shKDM2B-transduced MDA-MB-231 and MCF-10A cells were rescued with SLUG-WT, SLUG-4A, and SLUG-4D lentiviral constructs. Probing western blots of lysates derived from these cells with anti-HA or anti-Vinculin (loading control) antibodies showed that while the wild type and the SLUG-4A proteins were downregulated in shKDM2B-transduced cells, the phosphomimetic SLUG-4D mutant was not. **D.** shKDM2B-transduced MDA-MB-231 cells, engineered to express HA-tagged wild type SLUG (SLUG WT), SLUG-4A, or SLUG-4D, were treated with Cycloheximide (CHX) (30 μg/ml) and analyzed by immunoblotting at sequential time points for the abundance of the HA-tagged SLUG. Vinculin was the loading control. Given that shKDM2B downregulates SLUG, the loading in panels A and B was adjusted so that the amount of SLUG protein before the start of the treatment would be similar in the shControl and shKDM2B cells.

The homeostatic regulation of SLUG in normally growing cells, depends on GSK3-mediated phosphorylation at Ser92, Ser96, Ser100 and Ser104, which tags the protein for ubiquitination and proteasomal degradation ^11,36,37^. To determine whether the destabilization of SLUG by shKDM2B was due to stimulation of its homeostatic proteasomal degradation mechanism, we treated the EV and shKDM2B-tranduced MDA-MB-231 and MCF10A cells with the proteasome inhibitor Bortezomib (BORT) (0.5 μM) and we monitored the levels of SLUG over time, by western blotting. As expected, SLUG levels increased in Bortezomib-treated cells, but the increase was more pronounced in the control cells (Fig 2B), suggesting that the enhanced degradation in shKDM2B-transduced cells is proteasome-independent. To validate these data, we knocked down KDM2B in MDA-MB-231 and MCF10A cells transduced with constructs of wild type SLUG, or SLUG mutants in which all four phosphorylatable serine’s, were replaced by alanine (SLUG-4A), or aspartic acid (SLUG-4D). Measuring the abundance of SLUG by western blotting in these cells, showed that the knockdown of KDM2B promotes the downregulation of wild type SLUG and of the serine to alanine, but not the serine to aspartic acid mutant of SLUG (Fig 2C). In a parallel experiment, we monitored by western blotting the abundance of SLUG in CHX-treated (30 μg/ml), shKDM2B-transduced MDA-MB-231 cells, rescued with wild type or mutant SLUG, and we observed that whereas the wild type and the serine to alanine mutant of SLUG were degraded rapidly, the phosphomimetic serine to aspartic acid mutant was significantly more stable (Fig 2D).

The conclusion that the degradation of SLUG in shKDM2B-transduced cells is proteasome-independent, which was suggested by the preceding experiments, was further supported by the observation that shKDM2B promotes the phosphorylation of GSK3α/β at Ser21/Ser9 (Fig 3A). Phosphorylation at this site inhibits GSK3, whose kinase activity is required for the phosphorylation and proteasomal degradation of SLUG. To determine which GSK3 isoform undergoes phosphorylation in cells transduced with shKDM2B, we probed cell lysates of Control and shKDM2B-transduced MDA-MB-231 and MCF-10A cells with antibodies that distinguish phosphorylated GSK3α from phosphorylated GSK3β, and we observed that, although the phosphorylation of both isoforms was enhanced, the enhancement of GSK3β phosphorylation was more pronounced (Fig S2A).

**Figure 3:**
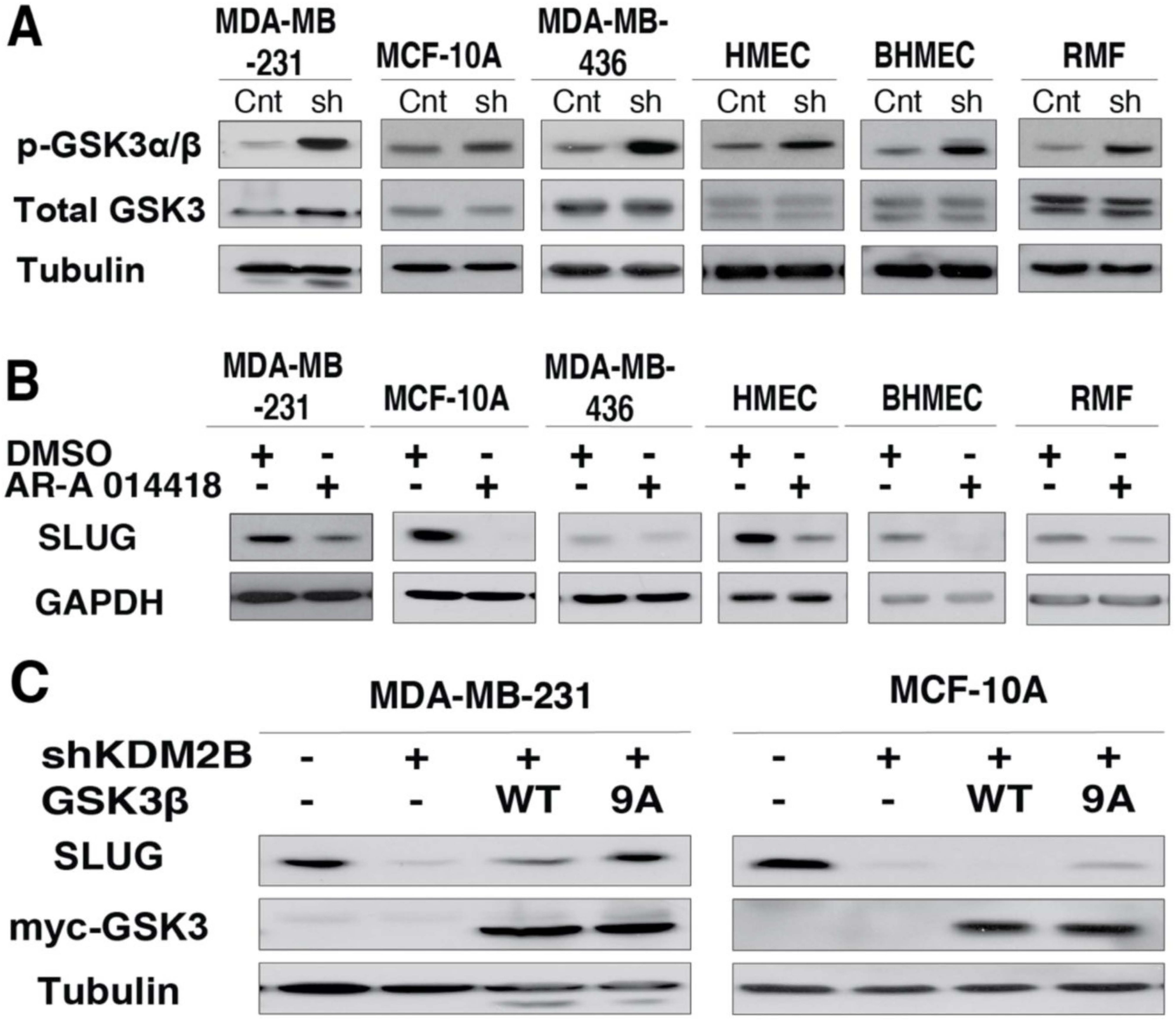
The knockdown of KDM2B destabilizes SLUG, in part by promoting the inhibitory phosphorylation of GSK3 at Ser9/Ser21. **A.** The six human mammary gland-derived cell lines in our panel, were transduced with shControl (Control) or shKDM2B (sh) lentiviral constructs. Probing immunoblots of cell lysates from these cell lines with anti-GSK3α/β anti-Total GSK3 and anti-Tubulin (loading control) antibodies, revealed that shKDM2B promotes GSK3 phosphorylation. **B.** The six human mammary gland-derived cell lines in our panel were treated with the GSK3 inhibitor AR-A 014418 (2 μM), or DMSO for 48 hours. Probing immunoblots of cell lysates from these cell lines with anti-SLUG or anti-GAPDH (loading control) antibodies, revealed that the inhibition of GSK3 results in the downregulation of SLUG. **C.** MDA-MB-231 cells (Left panel) and MCF-10A cells (Right panel), transduced with shControl or shKDM2B lentiviral constructs, were rescued with wild type GSK3β, or the constitutively active GSK3β S9A mutant. Probing immunoblots of lysates derived from these cells with the indicated antibodies, showed that the GSK3β S9A mutant rescues the SLUG downregulation fully in MDA-MB-231 cells and partially in MCF-10A cells.

### The shKDM2B-induced GSK3β inactivation contributes to the destabilization of SLUG

The results of the experiments in figures 2C, 2D and 3A suggested that the phosphorylation-induced inactivation of GSK3 may contribute to the destabilization of SLUG in KDM2B knockdown cells. To determine the validity of this hypothesis, we treated our cell line panel with the GSK3β inhibitor AR-A-014418 (2 μM) and we probed cell lysates harvested 48 hours later with the anti-SLUG antibody. The results in figure 3B provided support to the hypothesis by showing that, like the knockdown of KDM2B, AR-A-014418 also downregulates SLUG (Fig 3B). Additionally, the ectopic expression of both the wild type GSK3β and the constitutively active mutant GSK3βS9A, rescued the SLUG downregulation phenotype in shKDM2B-transduced cells, with the rescue by the GSK3βS9A mutant being more robust (Fig 3C). These data collectively support the hypothesis that the inactivation of GSK3 contributes to the destabilization of SLUG in KDM2B knockdown cells.

### Inhibition of the GSK3 catalytic activity promotes mesenchymal to epithelial transition (MET) and the differentiation of basal like breast cancer cell lines

SLUG expression promotes epithelial to mesenchymal transition (EMT) in MCF-10A cells ^11,38^. Given that inhibition of GSK3β induces a dramatic downregulation of SLUG, we asked whether chemical inhibitors of GSK3β promote mesenchymal to epithelial transition (MET), which is the reverse of EMT. In addition, we questioned whether treatment of MCF-10A cells with GSK3β inhibitors prevents the induction of EMT by TGFβ. To address the first question, we monitored the expression of SLUG, E-Cadherin and Vimentin in MCF-10A cells treated with AR-A-14418 (2 μM) for 7 days, and we observed that, although the SLUG levels returned to almost normal by day 4, after a precipitous drop by day 2, E-Cadherin levels gradually increased, while Vimentin levels gradually decreased (Fig S2B, Left panel). We conclude that GSK3β inhibition indeed promotes MET. To address the second question, we treated MCF-10A cells with TGFβ (4 ng/ml) for 7 days, in the presence or absence of AR-A-14418, and we observed that AR-A-14418 partially inhibits the induction of SLUG and Vimentin and promotes the upregulation, rather than the downregulation of E-Cadherin (Fig S2B, middle and right panels). We conclude that by interfering with the induction of SLUG, AR-A-14418 inhibits the induction of EMT by TGFβ.

Our earlier studies had shown that KDM2B promotes the self-renewal of cancer stem cells in basal-like breast cancer cell lines, and that the knockdown of KDM2B promotes differentiation toward a luminal phenotype ^11,22^. Given that AR-A-14418 also downregulates SLUG, we questioned whether GSK3β inhibition phenocopies the knockdown of KDM2B. To address this question, we treated MDA-MB-231 cells with AR-A-014418 (2 μM) or DMSO for 7 days, and we monitored them for the expression of the cancer stem cell markers CD44 and CD24 and the mammary epithelia differentiation markers CD49f and EpCAM, or CD49f and CD24. The results showed that AR-A-014418 treatment increases in the number of CD44low cells with a positive shift in CD24 expression (Fig S2C, left panel), suggesting a reduction in the abundance of cancer stem cells. Additionally, AR-A-014418 increased the number of CD49flow cells, with a positive shift in the expression of EpCAM or CD24 (Fig S2C, middle and right panels), suggesting differentiation from the basal-like to the luminal phenotype.

### GSK3β phosphorylation in shKDM2B-tranduced cell lines is under the control of complex phosphorylation and dephosphorylation-dependent mechanisms

GSK3β phosphorylation at Ser9 is regulated by kinases that phosphorylate this site and phosphatases that dephosphorylate it. Kinases known to phosphorylate this site include Akt, classical PKC family members, p70S6K, pp90RSK and SGK. To determine which of these kinases may be responsible for the basal and shKDM2B-induced GSK3β phosphorylation, we treated EV and shKDM2B-transduced MDA-MB-231 and MCF10A cells with AZD5363 (pan-Akt inhibitor) (0.5 μM) ^11,39^, R0-31-8220 (inhibitor of classical PKCs and PKCε) (0.1 μM) ^40^, PF-4708671 (p70S6K inhibitor) (10 μM) ^41^, or BI-D1870 (pp90RSK inhibitor) (10 μM) ^11,42^ Probing cell lysates harvested 2 hours later, with antibodies that recognize Ser9-phosphorylated or total GSK3β, showed that only the inhibitors of Akt and PKC inhibit the phosphorylation, excluding p70S6K and pp90RSK from the phosphorylation of GSK3β in these cells. However, these inhibitors reduced GSK3β phosphorylation not only in the KDM2B knockdown, but also in the control cells. (Fig S3A). Treatment of only the control cells with the SGK inhibitor GSK650394 (10 μM) ^43^ for 24 hours, showed that SGK may also contribute to GSK3β phosphorylation in these cells (Fig S3B).

A prerequisite for the kinase phosphorylating GSK3β in shKDM2B-transduced cells is that it is upregulated and/or activated, upon the knockdown of KDM2B. We therefore examined the expression and phosphorylation of Akt, the classical PKCs and SGK in control and shKDM2B-transduced cells. Figure S3C shows that the knockdown of KDM2B increased the phosphorylation of Akt in MDA-MB-231 and MCF-10A cells, but not in the remaining four cell lines, where Akt phosphorylation was decreased. It is interesting that the shKDM2B-induced changes in the phosphorylation of Akt, exhibited a perfect negative correlation with changes in the expression of INPP4B, a phosphoinositide phosphatase, which removes the D4 phosphate from the phosphorylated inositol ring of PtlIns3,4-P2 ^11,44,45^ (Fig S3D). This suggests that the differential regulation of Akt phosphorylation by shKDM2B in different cell lines may be due to the differential effects of shKDM2B on INPP4B expression. To determine the role of PKC isoforms on GSK3β phosphorylation in shKDM2B-transduced cells, we probed lysates of control and shKDM2B cells with a PKC phospho substrate antibody ^11,46^. The results revealed inhibition, rather than activation of PKC in all the shKDM2B-transduced cells (Fig S3E), excluding PKC as a candidate kinase for the shKDM2B-induced GSK3β phosphorylation. To determine the role of SGK isoforms in the shKDM2B-induced phosphorylation of GSK3β, we probed lysates of control and shKDM2B-transduced cells for SGK1, SGK2, SGK3 and phosphoSGK3 (Thr320) and we observed that the KDM2B knockdown results in the upregulation of SGK2 in MDA-MB-436, HMEC and RMF cells. SGK3 phosphorylation was also induced in MDA-MB-436 cells and HMECs (Fig S3F). We conclude that AKT, SGK2 and SGK3 may selectively contribute to GSK3 phosphorylation in different cell lines. The enhanced phosphorylation of SGK3 in MDA-MB-436 cells and HMECs could be explained by the increase in the abundance of INPP4B, which has been suggested to induce a signaling switch from AKT to SGK activation ^11,47^.

The potential role of phosphatase inactivation in the enhancement of GSK3β phosphorylation in shKDM2B-transduced cells, was addressed in MDA-MB-231 cells in which shKDM2B promotes GSK3β phosphorylation via AKT. Treatment of control and shKDM2B-transduced MDA-MB-231 cells with the pan-Akt inhibitor AZD-5363 (5 μM) and monitoring the GSK3β Ser9 phosphorylation for 30 minutes from the start of the treatment, revealed that the phosphorylation declines more rapidly in control than in shKDM2B-transduced cells (Fig S3G). These data suggest that the increase in GSK3 phosphorylation in KDM2B knockdown cells, is due at least in part, to the inactivation of phosphatases that dephosphorylate this site. Two phosphatases, PP1 and PP2A, are known to dephosphorylate GSK3 phosphorylated at Ser9/21 ^11,48^. Both phosphatases are complex multi-subunit metalloenzymes and their involvement in this process will be addressed in future studies.

### The knockdown of KDM2B in mammary cell lines destabilizes SLUG via calpain activation

To determine which protease family may be responsible for the destabilization of SLUG, we first treated EV control and shKDM2B-transduced MDA-MB-231 cells with increasing concentrations of chloroquine (CQ, 30 μM), 3-methyladenine (3-MA 10 mM), or concanamycin A (ConA 1 μM), which inhibit autophagy, by targeting different stages of the process ^11,49,50^. Monitoring the abundance of SLUG in control and shKDM2B cells treated with these inhibitors, confirmed that autophagy is not responsible for the shKDM2B-induced downregulation of SLUG, by showing that none stabilized it (Fig S4A). We next examined the role of calpains by monitoring the cellular levels of SLUG, before and after treatment of the same cells with a cell-permeable peptide derived from calpastatin (CAST 10 μM), an endogenous inhibitor of calpain-1, calpain-2 and calpain-9 ^11,51–53^. This experiment showed that SLUG was upregulated by calpastatin only in shKDM2B-transduced cells (Fig 4A), suggesting that calpastatin stabilizes SLUG selectively in these cells. This observation was interpreted to indicate that shKDM2B promotes the degradation of SLUG via calpain activation. Using an in vitro assay to measure calpain activity in cell lysates of control and shKDM2B-transduced cells, confirmed that the knockdown of KDM2B indeed promotes calpain activation (Fig 4B). The activation was reproducibly more robust in MCF10A than in MDA-MB-231 cells (Fig 4B), which was consistent with the finding that ectopic expression of GSK3βS9A rescues fully the shKDM2B-induced downregulation of SLUG in MDA-MB-231 and only partially in MCF10A cells (Fig 3C).

**Figure 4:**
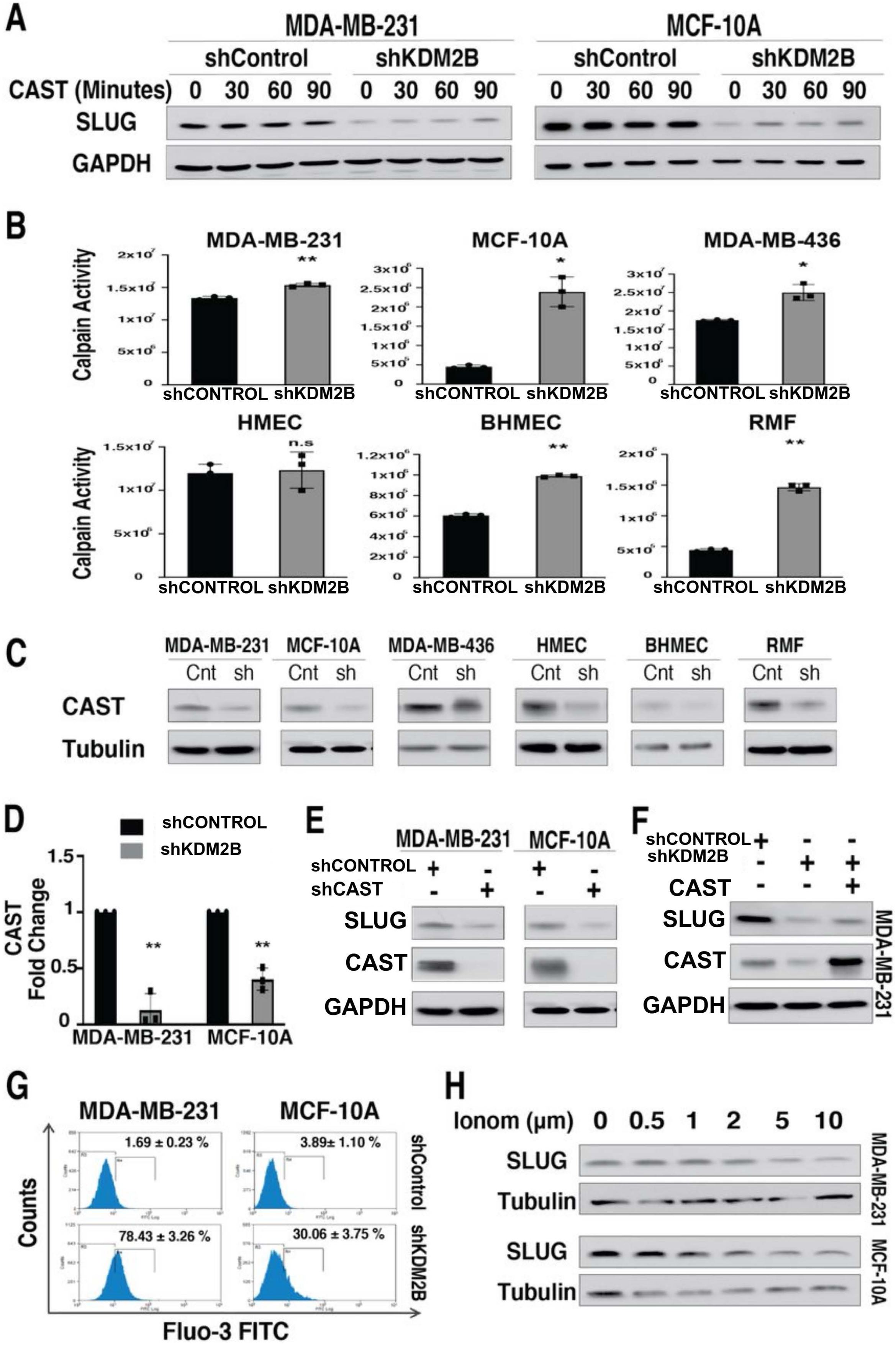
KDM2B knockdown promotes calpain-mediated degradation of SLUG: **A.** MDA-MB-231 cells (Left panel) and MCF-10A cells (Right panel), transduced with shControl or shKDM2B lentiviral constructs, were treated with a cell-permeable peptide derived from calpastatin (CAST), an inhibitor of calpains 1, 2, and 9 (10 μM). They were lysed and harvested at sequential time points, as indicated. Probing western blots of the harvested cell lysates with anti-SLUG or anti-GAPDH (loading control) antibodies showed upregulation of SLUG only in the shKDM2B-transduced cells, suggesting that the destabilization of SLUG by shKDM2B depends on the activation of calpains 1, 2, or 9.**B.** Measurement of the activity of calpains 1 and 2, using a luminescent assay in all six mammary cell lines in our panel, following transduction with shControl and shKDM2B constructs, confirmed the activation of these calpains by shKDM2B. Data are presented as mean ± SD of three independent experiments and asterisks indicate statistical significance (* p-value ≤ 0.05, ** p-value ≤ 0.01, *** p-value ≤ 0.001). **C.** Probing immunoblots of cell lysates from the cell lines in B with an anti-calpastatin antibody revealed a robust downregulation of calpastatin in all the shKDM2B-transduced cell lines. **D.** Measurement of the calpastatin mRNA levels by quantitative RT-PCR in shControl and shKDM2B-transduced MDA-MB-231 and MCF10A cells, revealed that calpastatin is downregulated by KDM2B at the RNA level. **E.** Probing immunoblots of shControl and shCAST (Calpastatin)-transduced MDA-MB-231 and MCF10A cells with the indicated antibodies, revealed that the knockdown of calpastatin is sufficient to downregulate SLUG. **F.** shKDM2B-transduced MDA-MB-231 cells were rescued with calpastatin and compare against shControl or shKDM2B cells. Probing immunoblots of lysates from these cells with the indicated antibodies, revealed that calpastatin rescues partially the downregulation of SLUG by shKDM2B. **G.** Fluo-3 staining and flow cytometry of the cells in D, revealed that shKDM2B increases the levels of intracellular Ca^2+^. **H.** MDA-MB-231 and MCF-10A cells were treated with ionomycin and they were lysed and harvested at 1 hour. Probing immunoblots of these lysates with anti-SLUG and anti-Tubulin (loading control) antibodies, revealed that SLUG is downregulated rapidly in response to ionomycin treatment.

To address the mechanism of calpain activation we first examined the expression of calpastatin and the calpastatin-inhibited calpains 1, 2 and 9 by western blotting, and we observed calpastatin downregulation in all the shKDM2B-transduced cell lines (Fig 4C) and calpain 1 and 2 upregulation in some (HMEC and RMF) (Fig S4B). Calpain 9 was downregulated rather than upregulated by shKDM2B in most cell lines, suggesting that it was not contributing to the phenotype. Since activated calpains degrade calpastatin ^11,54^, we also examined the abundance of calpastatin mRNA in control and shKDM2B cells, and we observed that it was also reduced by shKDM2B (Fig 4D and Fig S4C), suggesting that calpastatin downregulation precedes calpain activation. To determine whether the downregulation of calpastatin contributes to the shKDM2B-induced degradation of SLUG, we examined the abundance of SLUG in MDA-MB-231 and MCF-10A cells, before and after the knockdown of calpastatin, and we observed that calpastatin depletion results in SLUG downregulation in both cell lines (Fig 4E).Additionally, re-expression of calpastatin in shKDM2B-transduced MDA-MB-231 cells partially restored SLUG expression (Fig 4F).

The partial rescue of SLUG by calpastatin, in KDM2B knockdown cells, suggested that the calpastatin downregulation is one, but not the only factor responsible for the calpain-mediated SLUG degradation. Given that calpain is also activated by upregulation of the intracellular Ca^2+^ levels ^5^, we treated control and shKDM2B-transduced MDA-MB-231 and MCF10A cells with the calcium indicator Fluo-3 ^55^, and we measured intracellular Ca^2+^ by monitoring Fluo-3 fluorescence intensity via flowcytometry. The results confirmed that shKDM2B indeed upregulates intracellular Ca^2+^ (Fig 4G). To determine whether calpain activation induced by the upregulation of intracellular Ca^2+^ promotes SLUG degradation, we monitored the effects of increasing concentrations of ionomycin on calpain activity and SLUG abundance in MDA-MB-231 and MCF10A cells. The results confirmed both the activation of calpain (Fig S4D) and the degradation of SLUG by ionomycin (Fig 4H). We conclude that shKDM2B promotes calpain activation via a combination of calpastatin downregulation and intracellular Ca^2+^ upregulation.

### GSK3 inhibition and calpain activation synergize to destabilize SLUG

The preceding data raised the question whether GSK3 inhibition in shKDM2B-transduced cells activates calpain. To our surprise, although inhibition of GSK3 downregulates SLUG (Fig 3), it does not downregulate calpastatin and it does not activate calpain (Fig 5A and 5B). Based on these data, we hypothesized that the inhibition of GSK3 and the activation of calpain by shKDM2B, are two parallel processes which converge on SLUG, and that the inhibition of GSK3 may sensitize SLUG to degradation by the activated calpain (Fig 5C). To address this hypothesis, we treated MDA-MB-231 cells transduced with HA-tagged SLUG wild type, SLUG-4A and SLUG-4D, with ionomycin (1 hour), which activates calpains via KDM2B independent mechanisms. Probing lysates of these cells with an anti-HA antibody, revealed that wild type SLUG and SLUG4A, were significantly more sensitive to the treatment than SLUG4D (Fig 5D), suggesting that phosphorylation protects SLUG from calpain-mediated degradation. In a parallel experiment, HA-SLUG was immunoprecipitated from lysates of the same cells, and the immunoprecipitates were incubated with purified calpain for 30 minutes (Calpain-1 at 0.3 units/ml). Probing calpain-treated and untreated immunoprecipitates with an anti-HA antibody, revealed that whereas wild type SLUG and the 4A SLUG mutant were degraded, the 4D SLUG mutant was resistant to calpain-mediated degradation (Fig 5E), a finding consistent with the results of the experiment in figure 5D. In another experiment addressing the same question, recombinant wild type SLUG and the recombinant SLUG-4A mutant, were purified from *E Coli,* and they were used as substrates for GSK3β phosphorylation in an in vitro kinase reaction. GSK3β- and mock-phosphorylated wild type and mutant proteins were treated with purified calpain 1, as in the experiment in figure 5E. Monitoring the calpain-mediated degradation of these proteins by western blotting, revealed that phosphorylation protects wild type SLUG, but not its phosphorylation site mutant (Fig 5F). These results confirmed the proposed hypothesis, by showing that phosphorylation of SLUG by GSK3β partially protects it from degradation by calpain. Therefore, the inhibition of GSK3 via Ser21/Ser9 phosphorylation, renders SLUG more sensitive to calpain-mediated degradation (Fig 5C).

**Figure 5:**
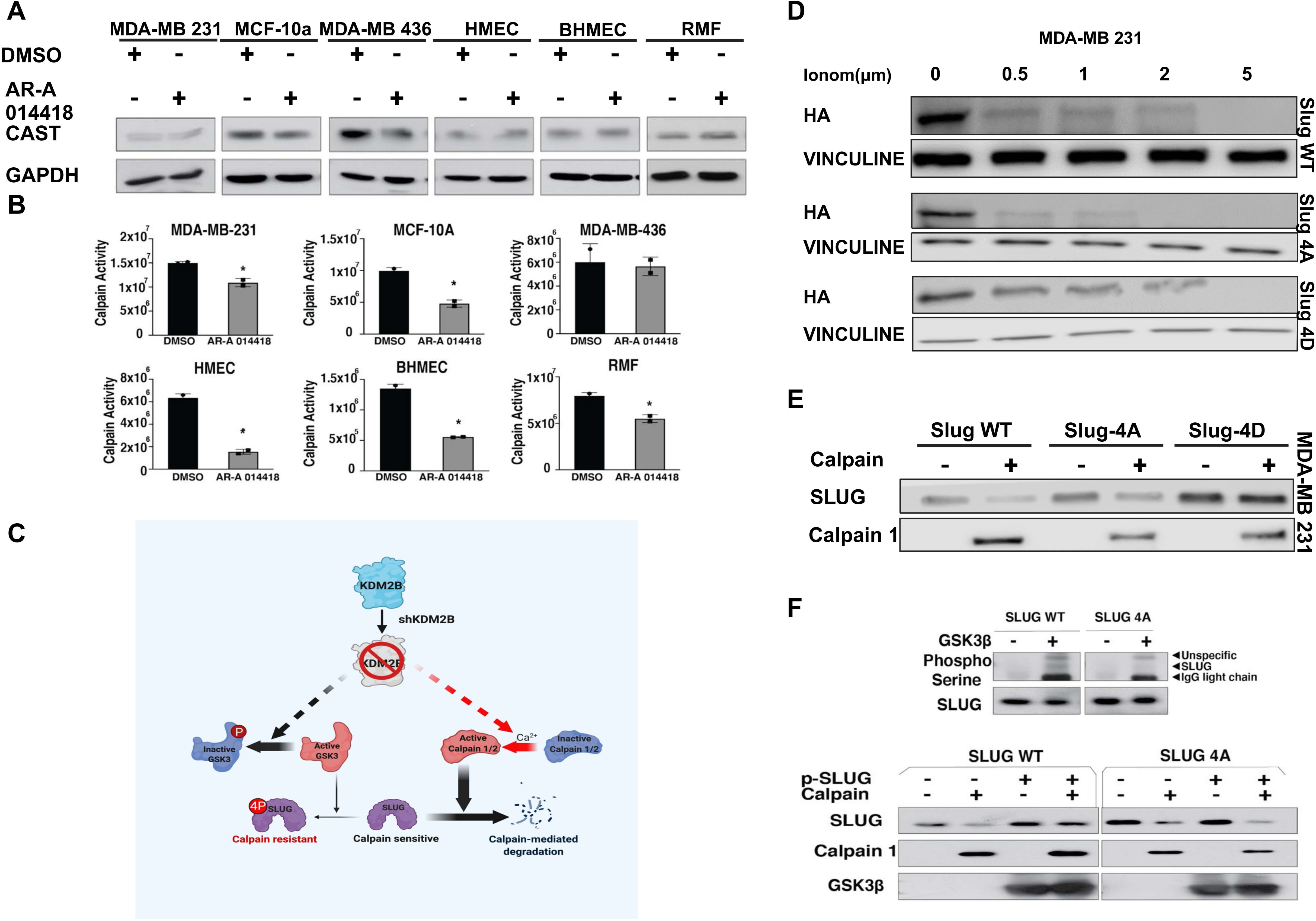
Phosphorylation by GSK3β protects SLUG from calpain-mediated degradation. **A.** The six human mammary gland-derived cell lines in our panel were treated with the GSK3 inhibitor AR-A 014418 (2 μM), or DMSO for 48 hours. Probing immunoblots of cell lysates from these cell lines with anti-CAST (calpastatin) or anti-GAPDH (loading control) antibodies, revealed that the inhibition of GSK3 results in weak downregulation of Calpastatin in some, but not all the cell lines. **B.** Measurement of the activity of calpains 1 and 2 by a luminescent assay in all six mammary cell lines in panel A, revealed that shKDM2B promotes the inhibition, rather than the activation of these calpains. Data are presented as mean ± SD of three independent experiments and asterisks indicate statistical significance (* p-value ≤ 0.05, ** p-value ≤ 0.01, *** p-value ≤ 0.001). **C.** The knockdown of KDM2B results in the inactivation of GSK3 and the activation of calpain. Both pathways converge on SLUG and promote its degradation. By inactivating GSK3, the knockdown of KDM2B blocks SLUG phosphorylation and renders SLUG sensitive to calpain-mediated cleavage. Therefore, the inactivation of GSK3 increases the calpain sensitivity of SLUG and acts synergistically with the activated calpain. **D.** MDA-MB-231 cells were treated with increasing concentrations of Ionomycin, which activates calpain via KDM2B-independent mechanisms. Probing western blots of cell lysates harvested 1 hour later, with anti-HA and anti-Vinculin (loading control) antibodies revealed that wild type SLUG and SLUG-4A were degraded more rapidly than the phosphomimetic SLUG-4D mutant. **E.** HA-tagged wild type SLUG, SLUG-4A and SLUG-4D, immunoprecipitated from MDA-MB-231 cells transduced with the corresponding SLUG constructs, were incubated with purified calpain 1 (0.3 units/ml) for 30 minutes. Western blotting of calpain-treated and untreated control immunoprecipitates, showed that whereas wild type SLUG and SLUG-4A were degraded by calpain, SLUG-4D was resistant to calpain-mediated degradation. **F.** *(Upper panel)* Wild type SLUG (SLUG-WT) and the mutant SLUG-4A were expressed and purified from E Coli. The purified proteins were phosphorylated by GSK3β in vitro. Probing an immunoblot of the GSK3β-phosphorylated and the mock-phosphorylated proteins with an anti-phospho-serine antibody confirmed the phosphorylation. *(Lower panel)* In Vitro degradation of phosphorylated and mock phosphorylated SLUG proteins (wild type and mutant) following treatment with purified calpain 1 (0.3 units/ml) for 30 minutes. Immunoblots of the products of the in vitro degradation reaction were probed with anti-SLUG, anti-calpain-1 and anti-GSK3β antibodies.

### The knockdown of KDM2B activates a paracrine mechanism, which results in GSK3 phosphorylation and calpain activation

The shKDM2B-induced GSK3, phosphorylation and cytoplasmic Ca2+ upregulation suggested the possibility that KDM2B knockdown activates a signaling cascade which ultimately results in the degradation of SLUG. To address this hypothesis, we probed lysates of control and KDM2B knockdown MDA-MB-231, MDA-MB-436 and HMEC cells with an anti-phosphotyrosine antibody (Millipore # 05-321). The results provided support to the hypothesis by showing that shKDM2B induces significant upregulation of tyrosine phosphorylation in all three cell lines (Fig 6A).

**Figure 6:**
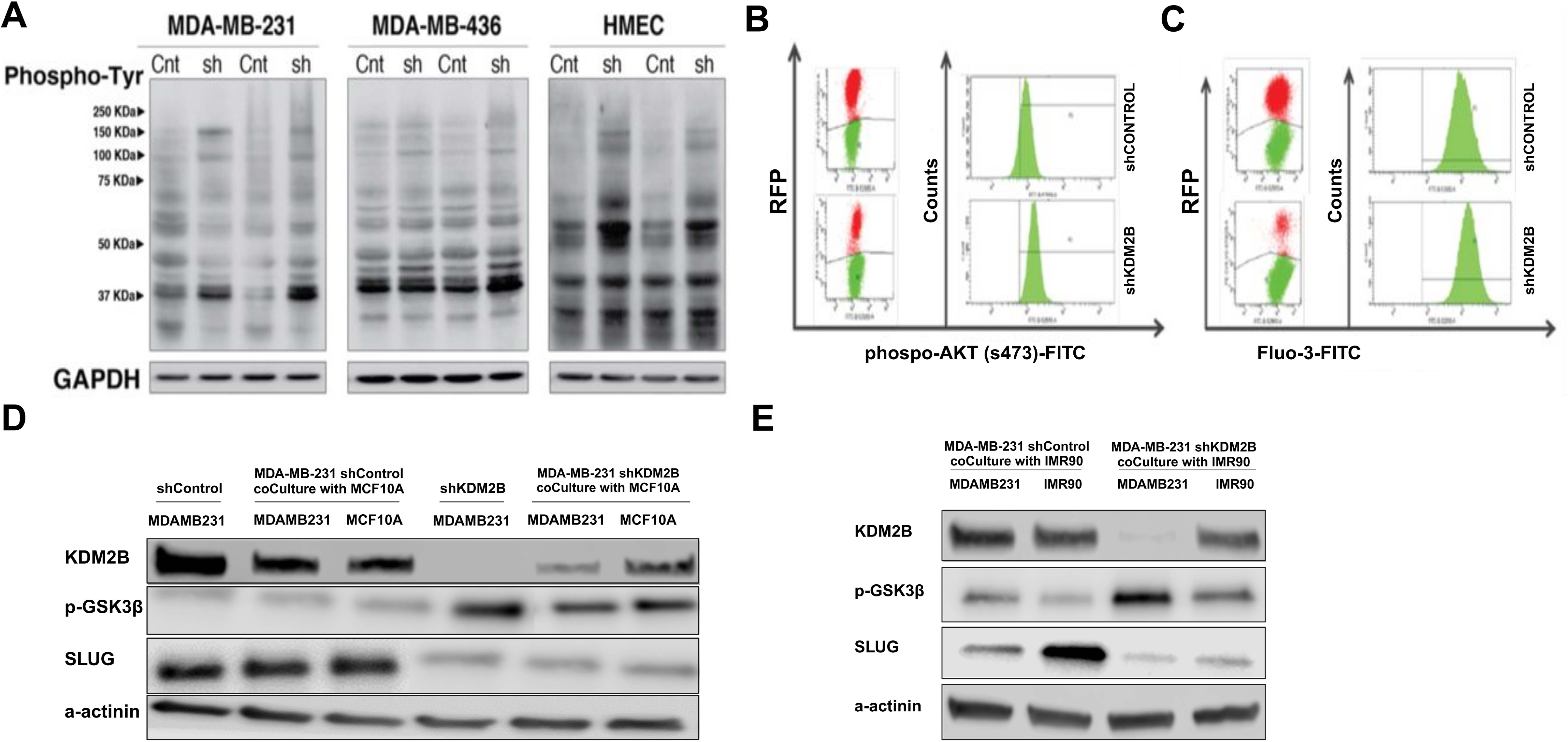
The knockdown of KDM2B in human mammary gland-derived cell lines activates GSK3β phosphorylation and calpain activation pathways via paracrine mechanisms. **A.** Probing immunoblots of MDA-MB-231, MDA-MB-436 and HMEC cells transduced with shControl or shKDM2B constructs with an antiphosphotyrosine antibody, revealed that shKDM2B promotes an increase in the abundance of tyrosine phosphorylated proteins. **B.** MDA-MB-231 cells engineered to stably express RFP, from a retrovirus construct, were transduced with shControl or shKDM2B constructs. The shControl and shKDM2B RFP-positive cells were co-cultivated with RFP-negative parental MDA-MB231 cells. Following co-cultivation, the cells were stained with an FITC-labeled anti phosphor-AKT (Ser473) antibody, and they were analyzed by flow-cytometry. **C.** Alternatively, the co-cultures were treated with the fluorescent Ca^2+^ indicator FLUO-3 AM and they were also analyzed by flow-cytometry. Gating on the RFP-negative cells revealed that they underwent shifts in the abundance of phosphor-AKT and Ca^2+^, when co-cultivated with shKDM2B cells, even though they were themselves, shKDM2B-negative. **D.** MDA-MB-231 cells, engineered to stably express GFP, were transduced with shControl or shKDM2B constructs. The shControl and shKDM2B GFP-positive cells were co-cultivated with GFP-negative MCF10A cells. Following a 48-hours co-cultivation period, the GFP-negative MCF10A cells were sorted from the GFP-positive MDA-MB-231 cells by flow-cytometry, and lysates of both, along with lysates of not co-cultivated control and shKDM2B MDA-MB231 cells, were probed with antibodies against KDM2B, phospho-GSK3β, SLUG, and α-Actinin (loading control). **E.** Control and shKDM2B-transduced GFP-positive MDA-MB-231 cells were co-cultivated with GFP-negative IMR-90 cells, and 48 hours later the GFP-positive and negative cells were sorted and probed with the same antibodies as in panel D.

The enhanced tyrosine phosphorylation of molecules in the range of ∼150 KD (Fig 6A), suggested that the knockdown of KDM2B activates a receptor tyrosine kinase (RTK)-dependent paracrine mechanism, which may ultimately lead to SLUG degradation. To address this hypothesis, we co-cultivated red fluorescent protein (RFP)-expressing MDA-MB-231cells transduced with shKDM2B, or the empty vector, with non-labelled parental MDA-MB-231 cells. Knocking down KDM2B in this cell line activates AKT by phosphorylation (Fig S3C). Staining the co-cultivated cells with an anti-phospho-Akt (phosphor-Ser473) antibody and analyzing them by flowcytometry, revealed that shKDM2B induces AKT phosphorylation not only in the shKDM2B-transduced/RFP-positive, but also in the co-cultivated//RFP-negative cells. (Fig 6B). In a parallel experiment, the same cells were treated with the Ca^2+^ sensor Fluo-3, and they were analyzed by flow-cytometry. This experiment revealed that intracellular Ca^2+^ was again increased in the cells co-cultivated with the shKDM2B-transduced cells (Fig 6C). The results of the preceding experiments confirmed that both AKT activation and Ca^2+^ influx, were induced via paracrine mechanisms.

To determine whether the paracrine mechanism activated by shKDM2B results in the phosphorylation of GSK3β, and the degradation of SLUG, we co-cultivated GFP-labeled EV and shKDM2B-transduced MDA-MB-231 cells, with unlabeled MCF10A or IMR-90 cells. Following 2 days of co-cultivation, GFP-negative MCF10A or IMR90 cells were sorted from the GFP-positive MDA-MB-231 cells, and lysates of both were probed with anti-phospho-GSK3β, and anti-SLUG antibodies. The results (Fig 6D, E) confirmed the hypothesis by showing that GSK3β undergoes phosphorylation, and that SLUG abundance is decreased in both the shKDM2B-transduced MDA-MB-231 cells, and their MCF10A and IMR90 neighbors.

To identify the RTKs activated by the shKDM2B-induced paracrine mechanism, we probed an anti-phosphotyrosine antibody array containing 228 antibodies (fullmoonbio.com), with lysates of empty vector control and shKDM2B MDA-MB-231 cells. The array included antibodies specific for tyrosine-phosphorylated RTKs, and for other proteins phosphorylated on tyrosine either directly by the RTKs, or by other tyrosine kinases in the pathways activated by these RTKs. To monitor the reproducibility of the results, all antibodies in the array were included in sextuplicate. The probing of the array was done by us, and the reading and analysis of the data were done by the array manufacturer. This screen identified 18 proteins, whose phosphorylation on tyrosine was increased in shKDM2B relative to the control cells. The top scoring proteins, based on their level of phosphorylation and the p-value, were the RTKs ERBB4 (HER4), FGFR1, TEK (TIE2), and MERTK, the cytoplasmic kinase PTK2B (PYK2/FAK2) and the tyrosine phosphorylated signaling protein STAT4 (Fig S5A and 7A).

To determine whether the signals transduced via tyrosine phosphorylation of these proteins are required for the shKDM2B-induced downregulation of SLUG, we examined whether their knockdown in MDA-MB-231 and MCF10A cells, rescues the shKDM2B-induced SLUG downregulation phenotype. The results showed that whereas the phenotype was rescued by the knockdown of two out of the four RTKs (ERBB4 and FGFR1) (Fig 7B), it was not affected by the knockdown of the other two (TEK and MERTK) (Fig S5B). The knockdown of the cytoplasmic tyrosine kinase PTK2B did not rescue the GSK3 phosphorylation, but rescued the SLUG downregulation (Fig 7C), while the knockdown of STAT4 did not rescue either (Fig S5C). The observation that both ERBB4 and FGFR1 are required for GSK3 phosphorylation and SLUG downregulation suggests that the two may function synergistically, as it has already been suggested in the prior literature ^56–59^. Because of the synergy, the depletion of either of the two may impact the activity of both.

**Figure 7:**
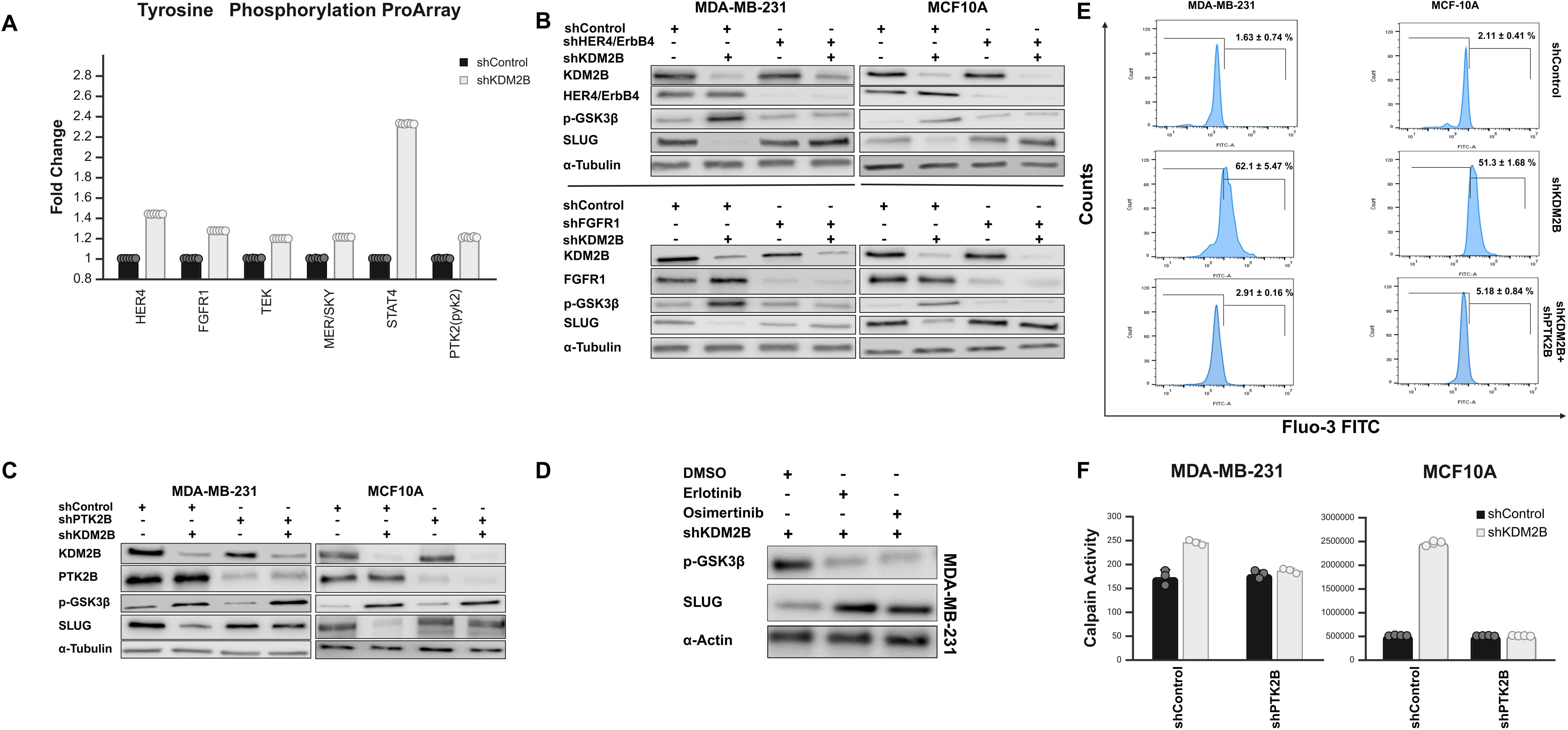
Tyrosine phosphorylation profiling revealed key mediators of the shKDM2B-induced paracrine signaling. **A.** Quantification of the shKDM2B-induced increase in the tyrosine phosphorylation of a set of proteins, which underwent robust increases in phosphorylation in shKDM2B-transduced MDA-MB-231 cells. Data were normalized with the level of phosphorylation in control cells being given the value of 1. Bars show the mean value of phosphorylation relative to the control. Individual data points are also shown, and they are very tightly clustered, confirming the reproducibility of the results. **B.** The knockdown of ERBB4 and FGFR1 rescues the GSK3 phosphorylation and SLUG degradation phenotype of the KDM2B knockdown. Western blots of lysates of control and KDM2B knockdown MDA-MB-231 and MCF10A cells were transduced with constructs of shERBB4, shFGFR1 or the shControl, and they were probed with the indicated antibodies. **C.** The knockdown of PTK2B rescues partially the SLUG degradation phenotype, but not the GSK3 phosphorylation phenotype of the KDM2B knockdown. Western blots of lysates of control and KDM2B knockdown MDA-MB-231 and MCF10A cells were transduced with an shPTK2B construct, or with the shControl, and they were probed with the indicated antibodies. **D.** The EGFR inhibitors Erlotinib and Osimertinib rescue the GSK3 phosphorylation and SLUG degradation phenotype of the KDM2B knockdown. Western blots of lysates of KDM2B knockdown MDA-MB-231 cells were treated with Erlotinib (7 μM), Osimertinib (5 μM), or DMSO for 24hours, and they were probed with the indicated antibodies. **E.** The knockdown of PTK2B rescues the Ca2+ upregulation phenotype of the KDM2B knockdown. Control and KDM2B knockdown MDA-MB-231 and MCF10A cells were transduced with an shPTK2B construct. Intracellular calcium levels in these cells were measured by flowcytometry, following treatment with the fluorescent Ca^2+^ indicator FLUO-3 AM. **F.** Measurement of the activity of calpains 1 and 2, in control KDM2B knockdown MDA-MB-231 and MCF10A cells, transduced with an shPTK2B construct, or the shControl. Measurements were performed with the luminescent assay described in the methods section. Data are presented as mean ± SD of three independent experiments and asterisks indicate statistical significance (* p-value ≤ 0.05, ** p-value ≤ 0.01, *** p-value ≤ 0.001).

The EGFR family has four members, which following ligand binding, are activated by forming homodimers, or heterodimers. At least eleven ligands are known to bind members of the EGFR family, with some of them binding selectively one, and others binding multiple members. One of the EGFR family members (ERBB2/HER2) has no known ligands, and another one (ERBB3) has a defective kinase domain. These members function by dimerizing with other members. ERBB4, the family member whose phosphorylation was linked to the knockdown of KDM2B, is known to bind three ligands, HB-EGF, EPR (EREG) and BTC, the first two of which were upregulated in shKDM2B cells (Fig S6A). These ligands also bind EGFR, which heterodimerizes with ERBB4. EGFR is the only member of the family, recognized by an additional set of four ligands, EGF, TGF-α, ARG (AREG) and EGN (EPGN), which should also activate ERBB4, because of the dimerization of ERBB4 with EGFR^60^. Importantly, based on our RNA-Sec data, the first three of these ligands were also upregulated in shKDM2B cells (Fig S6A). Based on the preceding data, we hypothesized that inhibition of EGFR with small molecule inhibitors, would also rescue the GSK3 phosphorylation and SLUG degradation phenotype in shKDM2B cells. Monitoring GSK3β phosphorylation and SLUG abundance by western blotting in shKDM2B cells treated with the EGFR inhibitors Osimertinib or Erlotinib confirmed the hypothesis (Fig 7D). We should add here that EGFR family members are known to be promiscuous, dimerizing with members of other RTK families, which suggests that inhibitors of other RTKs ^57^ may also rescue the SLUG degradation phenotype in shKDM2B cells. To determine whether the activation of EGFR family members in KDM2B knockdown cells was relevant to human cancer, we analyzed the reverse phase proteomic array (RPPA) data of the 1014 cases of breast cancer in the 2019 Firehose database (TCGA), for the expression and phosphorylation of the members of this RTK family. The results revealed significantly higher levels of ErbB2, ErbB2_pY1248, and ErbB3 in tumors with low levels of KDM2B (Fig S6B), suggesting that the paracrine mechanism activated by the knockdown of KDM2B in cultured cells, may indeed be relevant to human breast cancer.

The experiment in figure 7C, showing that the knockdown of PTK2B rescues the shKDM2B-induced SLUG degradation, but not the GSK3 phosphorylation phenotype, identifies a link, connecting the ERBB4-dependent paracrine mechanism with Ca^2+^ influx, one of the arms of the SLUG degradation pathway. PTK2B (PYK2), functions both as a sensor and effector of Ca++ influx. The KFL linker in PTK2B interacts with Ca^2+-^ bound Calmodulin (CaM). This results in PTK2B self-association, autophosphorylation and the downstream modulation of the activity of Ca^2+^ channels ^61,62^. Following its activation by RTK-initiated signals in KDM2B knockdown cells therefore, PTK2B contributes to SLUG degradation, by upregulating cytoplasmic Ca^2+^ levels and inducing calpain activation (Fig 7E and 7F).

The store-operated Ca^2+^ channels, ORAI3 and ORAI1 may also contribute to the Ca^2+^ upregulation in shKDM2B cells. First, our RNA-seq studies revealed significant upregulation of ORAI3 in KDM2B knockdown MDA-MB-231 cells (2.28-fold increase, FDR=0.00055). Consistent with this observation, the cBioportal-analyzed TCGA data on breast cancer revealed a negative correlation between the two (Spearman correlation coefficient −0.34) (https://www.cbioportal.org). The expression of ORAI1^63^ was not significantly affected in KDM2B knockdown cells. However, the gene encoding ORAI1 maps immediately upstream of the gene encoding KDM2B, raising the possibility that the two may be coregulated. Strong support to this hypothesis was provided by the observation that the expression of KDM2B and ORAI1 exhibit a positive correlation in human breast cancer (Spearman correlation coefficient 0.38) (https://www.cbioportal.org), suggesting that a compensatory increase in the transcription of KDM2B, which may occur in KDM2B knockdown cells, is likely to upregulate both KDM2B and ORAI1.

However, analysis of the reverse phase proteomic array (RPPA) data of the 1014 cases of breast cancer, in the 2019 Firehose TCGA database, revealed significantly higher levels of ErbB2, ErbB2_pY1248, and ErbB3 in tumors with low levels of KDM2B (Fig S4). These data suggest that the knockdown of KDM2B may result in the upregulation and activation of ErbB2 and ErbB3. The receptor tyrosine kinases and the tyrosine phosphorylation pathways activated by the knockdown of KDM2B, will be addressed in future studies.

## DISCUSSION

Our earlier studies had shown that KDM2B, in concert with EZH2, represses the expression of a set of microRNAs which target multiple components of the polycomb complexes PRC1 and PRC2. As a result, KDM2B upregulates PRC1 and PRC2 and promotes the self-renewal of breast cancer stem cells. As expected from these data, the knockdown of KDM2B depleted breast cancer cell lines of cancer stem cells, by promoting their differentiation in culture and in xenografts growing in orthotopically transplanted immunodeficient mice ^22^. Given that other studies had shown that the self-renewal and differentiation of mammary epithelial stem cells also depends on SLUG, SNAIL and SOX9 ^11,33–35^, we hypothesized that KDM2B may also regulate the expression of these molecules.

Data presented in this report confirmed this hypothesis, by showing that shKDM2B downregulates these molecules, and that their downregulation is posttranscriptional. Additionally, they showed that the downregulation of at least one of these molecules (SLUG), is due to destabilization and degradation. The potential interdependence of the regulation of PRC complexes and SLUG/SNAIL/SOX9 by KDM2B, will be addressed in future studies.

Earlier studies had shown that the abundance of SLUG in normally growing cells is under the homeostatic control of the proteasome. SLUG is normally phosphorylated at Se92, Ser96, Ser100 and Ser104 by GSK3, which tags SLUG for ubiquitination and proteasomal degradation ^11,36,55,64^. Given the data in this report, showing that the knockdown of KDM2B downregulates SLUG by promoting its degradation, we examined the expression and phosphorylation of GSK3, and to our surprise, we observed that shKDM2B promotes GSK3 inactivation via phosphorylation at Ser21/Ser9. This observation, combined with the results of additional pharmacologic and genetic studies, confirmed that the shKDM2B-induced SLUG degradation does not depend on the proteasome, but on calpain activation, which is due to the upregulation of intracellular Ca^2+^ and the downregulation of calpastatin. Given that calpastatin selectively inhibits calpains 1, 2 and 9 ^11,51–53^, this finding limited the spectrum of calpains that may be responsible for SLUG degradation by shKDM2B to calpains 1, 2 and 9. Of these calpains, the expression of calpain 1 and calpain 2 remains unchanged, or increases in shKDM2B-transduced cells, while the expression of calpain 9 decreases in most of the cell lines we tested. We conclude that SLUG degradation is mediated primarily by calpains 1 and 2.

Our data showing that the knockdown of KDM2B promotes the degradation of SLUG via GSK3 inactivation and calpain1/2 activation raised the question whether the activation of calpains 1 and 2 depends on the inhibition of GSK3. To our surprise, although inhibition of GSK3 downregulated SLUG, it did not activate calpains 1 and 2. This led us to conclude that GSK3 inhibition and calpain activation are parallel processes, which however may converge on SLUG, and that the inhibition of GSK3 may sensitize SLUG to calpains, by preventing SLUG phosphorylation. Importantly, there is precedence for the inhibition of calpain-mediated proteolysis of target proteins by phosphorylation ^11,65–68^ and one of the proteins whose stability is regulated by this mechanism is GSK3 itself ^11,67^. Data presented in this report confirmed that calpain-dependent SLUG degradation is indeed inhibited by GSK3-mediated phosphorylation. The preceding data show that the phosphorylation of SLUG by GSK3, which is required for the homeostatic degradation of SLUG by the proteasome, may also functionally protect SLUG from fluctuations of calpain activity that may occur under physiological conditions, during normal cell growth. Given the profound effects of SLUG downregulation in cellular physiology, we propose that the elegant switch mechanism regulated by the coupling of the two SLUG degradation mechanisms was designed to functionally stabilize SLUG, while maintaining it in a state poised for rapid degradation, in response to differentiation signals. This hypothesis is supported by our data showing that pharmacological inhibition of GSK3 downregulates SLUG and prevents EMT in cells treated with TGF-β. The ability of GSK3 blockade to downregulate SLUG and drive mammary cell differentiation ^11,69,70^, suggests that in the absence of GSK3-mediated phosphorylation, SLUG may be sensitized to basal calpain activity levels in cancer cells, or to other currently unknown proteases. Additionally, it suggests that GSK3 inhibition may be a promising strategy for the treatment of basal-like breast cancer. GSK3 inhibitors are available, and at least one of them (9-ING-41) is currently being tested in therapeutic trials focusing on different types of human cancer (https://clinicaltrials.gov/).

The relative contribution of GSK3 inactivation and calpain activation in the degradation of SLUG by shKDM2B may vary between cell lines. This is suggested by the comparison of the magnitude of calpain activation in shKDM2B-transduced MDA-MB-231 and MCF10A cells and the ability of the constitutively active mutant GSK3β-S9A to rescue the calpain-mediated cleavage of SLUG in the two cell lines. Thus, in MDA-MB-231 cells in which calpain activation was weak (Fig 5B), the rescue by GSK3β-S9A was complete (Fig 3C Left panel), while in MCF10A cells, in which calpain activation was strong (Fig 5B), the rescue was partial (Fig 3C Right panel).

Treatment of MDA-MB-231 and MCF10A cells with several kinase inhibitors revealed that under steady state conditions, GSK3β is phosphorylated at Ser9, primarily by AKT, PKC and SGK ^11,48^. However, the enhancement of GSK3 phosphorylation in shKDM2B-transduced cells may be caused primarily by inhibition of its dephosphorylation, with contributions from AKT and SGK, which are activated selectively downstream of shKDM2B in different cell lines. AKT for example may contribute to the increase in GSK3 phosphorylation in shKDM2B-transduced MDAMB-231 and MCF-10A cells but not in the other cell lines, probably because of the differential regulation of INPP4B by KDM2B in these cells. Analysis of the breast cancer CPTAC dataset ^11,71^ for correlations between GSK3 phosphorylation at Ser9/21 and the abundance of different PP1 and PP2A subunits (total and phosphorylated at various sites), provided evidence suggesting regulatory mechanisms that may be KDM2B-dependent and may contribute to the abundance of GSK3 phosphorylation in human breast cancer (See Tables S1 and S2). The upregulation of intracellular Ca^2+^ in KDM2B knockdown cells depends on PTK2B, which is a downstream target of the paracrine mechanism described in this report. Additional contributors could be the store operated channels ORAI3, and perhaps ORAI1, whose expression may be regulated by KDM2B. Whether PTK2B and the Ca^2+^ channels operate in the same or parallel pathways, has not been addressed.

Experiments addressing the important question of how the knockdown of a histone demethylase induces GSK3 phosphorylation and calpain activation, revealed that both events depend on shKDM2B-induced paracrine signaling. Specifically, knocking down KDM2B in RFP-expressing MDA-MB-231 cells co-cultivated with parental RFP-negative MDA-MB-231 cells, resulted in AKT phosphorylation and Ca^2+^ influx, not only in the RFP-positive cells in which KDM2B was knocked down, but also in the RFP-negative parental cells. Additionally, the co-cultivation of shKDM2B-transduced GFP-tagged MDA-MB-231 cells with MCF10A or IMR90 cells, resulted in GSK3 phosphorylation and SLUG downregulation in both the KDM2B knockdown cancer cells, and their neighbors. The shKDM2B-induced activation of paracrine signaling suggested by these experiments, was supported by studies showing that the knockdown of KDM2B upregulates tyrosine phosphorylation in all tested cell lines. A screen for proteins whose tyrosine phosphorylation was enhanced in shKDM2B cells, identified FGFR1, EGFR family members and PTK2B, as top candidates for the transduction of the shKDM2B-induced paracrine signals. Further studies confirmed that FGFR1 and EGFR family members initiate the signaling cascades and PTK2B promotes the upregulation of intracellular Ca^2+^, a signal for calpain activation ^5^. Analysis of the transcriptomes of control and shKDM2B-transduced MDA-MB-231 cells, identified growth factors that are upregulated by shKDM2B, and may be responsible for the activation of EGFR family members. They also identified additional growth factors that are regulated by KDM2B, and they may contribute to the shKDM2B-induced paracrine signaling. The negative correlation between KDM2B and levels of ErbB2, ErbB2_pY1248, and ErbB3 in human breast cancer, supports the significance of the paracrine mechanism described in this report in human cancer.

In summary, data presented in this report identify a novel tyrosine kinase receptor-dependent paracrine mechanism (Fig S7), which is activated by KDM2B depletion and promotes the calpain-mediated degradation of SLUG and the differentiation of mammary epithelia and basal-like breast cancer cells. KDM2B knockdown destabilized SLUG through convergence of calpain activation, and GSK3 inactivation. The impact of this paracrine mechanism on the tumor microenvironment remains to be determined. The coupling of the shKDM2B-induced calpain mediated SLUG degradation, which is inhibited by GSK3-mediated SLUG phosphorylation, with the GSK3 phosphorylation-dependent homeostatic proteasomal degradation of SLUG, suggests that GSK3 may protect SLUG from degradation that may be induced by small fluctuations in calpain activity in normally growing cells.

## MATERIAL AND METHODS

### (For details, see “Supplementary Materials and Methods”)

#### Cell culture

Human mammary gland derived cell lines and culture media listed are listed in Table S3. Cells were transduced with packaged lentivirus or γ-retrovirus constructs, using standard procedures. Inhibitors, growth factors and chemicals used for cell culture experiments are detailed in Table S4. Treatment details are provided in the results section.

#### shRNAs and expression vectors. Cloning and site-directed mutagenesis

Lentiviral shRNA Constructs are listed also in Table S9.

Expression constructs and their generation are described in Supplementary Material and Methods.

DNA primers for subcloning and site-directed mutagenesis are listed in Table S5. Gene blocks used for the construction of SLUG mutants are listed in Table S6.

#### RNA-Seq and Ribosome Profiling

Transcriptomic and ribosome profiling data in this report were derived from the analysis of previously reported RNA-seq and Ribo-Seq studies in control and KDM2B knockdown MDA-MB-231 cells, ^72,73^.

#### Real time RT-PCR

cDNA was synthesized using the Retroscript reverse transcription kit (Thermo Fisher Scientific, Cat. #AM1710) and total cell RNA extracted using Tripure (Sigma-Aldrich Cat. #11667157001). Gene expression was quantified by real time RT-PCR, using SYBR Green I master mix (Roche, Cat. #04887352001). The primer sets used for all the real time PCR assays are listed on the Table S7.

#### Immunoprecipitation and Immunoblotting

Cells were lysed in Triton X-100 lysis buffer (20 mM Tris (pH 7.5), 150 mM NaCl, 1 mM EDTA, 1 mM EGTA, 1% Triton X-100) or RIPA buffer (Thermo Scientific, cat. no 89900) supplemented with protease and phosphatase inhibitor cocktails (Sigma-Aldrich, Cat. # 11697498001 and Cat. # 4906837001 respectively). Immunoblotting of Triton X-100 and RIPA lysates and immunoprecipitation of Triton X-100 lysates followed by immunoblotting, were carried out using standard procedures. All antibodies used in the experiments in this report are listed in Table S8.

#### Intracellular calcium levels

Cells loaded with Fluo3-AM (Molecular probes, Cat. #F1241) were trypsinized and the intensity of fluorescence induced by Ca^2+^ binding was measured by flow cytometry.

#### Calpain activity assay

Cells were lysed in an EDTA-free lysis buffer containing 20 mM Tris-HCl pH=7.5 and supplemented with a protease inhibitor cocktail (Sigma-Aldrich, Cat. # 4693159001). Calpain activity was measured using the Calpain-Glo assay (Promega, Cat. # G8501)

#### Calpain-mediated cleavage of SLUG in vitro

3xFlag-SLUG-WT and 3xFlag-SLUG-4A proteins were immunoprecipitated from cells transduced with the corresponding constructs and the immunoprecipitates were phosphorylated by GSK3β in vitro. Following confirmation of the selective phosphorylation of the immunoprecipitated proteins, wild type and mutant proteins were treated similarly with calpain 1 (Sigma-Aldrich, Cat. #C6108) in vitro and their cleavage was assessed by immunoblotting. pInd20 constructs expressing HA-SLUG-WT, HA-SLUG-4A, and HA-SLUG-4D were transduced into MDA-MB-231 cells, and expression was induced with 1 μg/ml doxycycline. Forty-eight hours later, wild type SLUG and the SLUG mutants were immunoprecipitated from the cleared lysates of these cells with anti-HA antibodies, and the immunoprecipitates were incubated with calpain 1 (Sigma-Aldrich, Cat. #C6108) for 30 minutes. SLUG cleavage was assessed by immunoblotting.

#### Flow cytometry

Following trypsinization, 500,000 cells were stained with antibodies specific for phosphor-AKT (Ser473), or cell surface differentiation markers, using standard procedures, and the intensity of staining was measured by flowcytometry.

#### Statistical analysis

All experiments were performed in triplicate, and data were presented as the mean ± SD. Statistical analyses were tailored to the specific experimental design: comparisons between two groups were conducted using either the student’s t-test or the Mann–Whitney U-test, depending on sample distribution. For broader analyses, multiple unpaired t-tests were performed with Holm-Šídák correction for multiple hypothesis testing. Statistical analyses were conducted using GraphPad Prism and SPSS version 19.0. p-values and adjusted p-values of less than 0.05 were considered significant.

## Data Access Statement

All omics data referred to in this manuscript were included in prior publications^72^ and they are already publicly available.

## ACKOWLEDGMENTS

The authors wish to thank Dr Philip Hinds and Charlotte Kuperwasser (Tufts University School of Medicine) and all the members of the Tsichlis lab for helpful discussions. We also wish to thank Dr Charlotte Kuperwasser (Tufts University School of Medicine) for kindly providing us with cell lines used in this research. Finally, we wish to thank Dr Joseph Amann (The Ohio State University) for providing us with samples of Osimertinib and Erlotinib. This work was supported by the NIH grant R01 CA109747 and by start-up funds from the OSUCCC to P.N. Tsichlis. Elia Aguado Fraile was supported by a postdoctoral fellowship from the Alfonso Martin Escudero Foundation (Spain). All the Figures including the graphical abstract and the schematic diagrams were created with BioRender.com.

## Author Contributions

E.A.F. and B.S. contributed to the design of study and performed key experiments, analyzed data, and wrote parts of the manuscript. V.A. and Z.T.S. performed experiments and contributed to data interpretation. O.S. and E. C. contributed to experimental design and some of the reported experiments. M.D.P. performed data analysis. P.N.T. conceived the project, supervised research, and edited the manuscript. All authors reviewed and approved the final manuscript.

## Ethics Statement

Human or animals were not direct subjects in any of the experiments in this study.

