## Supplementary information for "KDM2B Silencing Elicits a Paracrine Mechanism Which Destabilizes SLUG, Promoting Differentiation of Basal-Like Breast Cancer Cells"

### SUPPLEMENTARY MATERIAL AND METHODS

#### Cell Culture

Human mammary gland-derived cell lines were cultured in the media listed in Table S3. Immortalized human mammary cell lines HMEC, BHMEC, and RMF were kindly provided by Dr. Charlotte Kuperwasser [74,75]. Retroviral constructs were packaged in 293T cells by transient co-transfection with an amphotropic packaging construct (Ampho-pac). Lentiviral pLK0.1, pLenti-CMV-DEST, and Pin20 constructs were packaged in 293T cells by transient co-transfection with psPAX2 and pMD2.G. Transfections were carried out using X-tremeGENE 9 DNA Transfection Reagent (Sigma-Aldrich, Cat. #6365787001) or Lipofectamine 3000 (Thermo Fisher Scientific, Cat. #L3000001) according to the manufacturer's instructions.

Infections of cancer cell lines were carried out in the presence of 5 μg/ml polybrene (Sigma-Aldrich, Cat. #107689). Immortalized mammary cell lines were pre-treated for 45 minutes with 25 μg/ml DEAE dextran (Sigma-Aldrich, Cat. #D9885) and subsequently infected without polybrene. Forty-eight hours after exposure to the virus, cells were sorted for GFP or selected with puromycin (2 μg/ml), hygromycin (300 μg/ml), blasticidin (5 μg/ml), or neomycin (500 μg/ml) depending on the selection marker of the vector. After neomycin selection, cells expressing SLUG-WT, SLUG-4A, or SLUG-4D constructs in pInd20 were induced with 1 μg/ml doxycycline for 48 hours. Cells transduced with pInd20 constructs were cultured in media supplemented with Tet-free FBS (Omega Scientific Inc., Catalog No. NC0290780), prior to the induction with doxycycline. Inhibitors, growth factors, and chemicals used for cell culture experiments are detailed in Table S4.

#### shRNAs and Expression Vectors. Cloning and Site-directed Mutagenesis

Lentiviral shRNA constructs for human KDM2B were purchased from Dharmacon (Cat. #RHS3979-201837202, clone ID: TRCN0000118437) [13]. An additional KDM2B shRNA construct, pLKO.1-shKDM2B (shRNA TRCN0000238779) was purchased from Millipore-Sigma. A set of five lentiviral shRNA clones for calpastatin were also purchased from Dharmacon (RHS4533-EG831). Out of the five, clone TRCN0000073638 induced the most robust knockdown, as determined by western blotting. Lentiviral shRNA constructs targeting several additional genes were obtained from Millipore

Sigma. These included shRNAs for STAT4 (Cat. #TRCN0000020894 and Cat. #TRCN0000020895), FGFR1 (Cat. #TRCN0000312574), TEK (Cat. #TRCN0000000412), PTK2B (Cat. #TRCN0000231523), Erbb4/HER4 (Cat. #TRCN0000001411), and MERTK (Cat. #TRCN0000000865).

**shRNA Constructs are listed also in Table S9.**

Transient expression constructs 3xFLAG-SLUG-WT and 3xFLAG-SLUG-4A (S92A / S96A / S100A / S104A) were a kind gift from Dr. Stephen J. Weiss [76]. FLAG-SLUG-WT and FLAG-SLUG-4A cDNAs were subcloned into pBabe-Hygro, using primers listed in Table S5. An additional set of SLUG constructs (HA-SLUG-WT, HA-SLUG-4A, and HA-SLUG-4D) in the Tetracycline-inducible lentiviral vector pInd20 [77], were generated by us from pENTR-SLUG-WT by exchanging wild type sequences with mutant gene blocks (IDT) (Supplementary Table S7) via reactions catalyzed by the NEB HiFi DNA Assembly Master Mix (New England BioLabs, Cat. #E2621S). These constructs were validated through full plasmid sequencing (Plasmidsaurus) before recombination into the Pin20 lentiviral vector using Gateway LR clonase II reactions (Thermo Fisher Scientific, Cat. #11791020). The final Pind20 constructs were verified again by Plasmidsaurus and subsequently packaged into lentivirus using psPAX2 and pMD2.G in Lenti-X 293T cells.

**Mutagenesis primers and primers for insertion mutant gene blocks are listed also in Table S5.**

GSK3β (NM_002093.3) with a C-terminal Myc tag was initially cloned into pBABE-Puro. Subsequently, GSK3β-myc was transferred into pENTR/D-Topo and recombined into pLenti-CMV-DEST using the gateway cloning system. Primers used for subsequent subcloning steps are listed in Table S5. GSK3β-S9A-myc was generated by site-directed mutagenesis using the QuikChange II kit (Agilent Cat. #200521).

KDM2B WT and point and deletion mutants had been described previously [7]. KDM2B cDNA was transferred from retroviral vectors into pENTR and subsequently recombined into pLX304, using the Gateway LR clonase II system (Thermo Fisher Scientific Cat. # 11791020). Calpastatin cDNA in pDONR221 was purchased from DNAsu (Cat # HsCD00040354) and transferred to pLX304 as previously described.

**Real time RT-PCR**

Total cell RNA was extracted using Tripure (Sigma-Aldrich Cat. #11667157001). cDNA was synthesized from 1.0 µg of total RNA, using random-decamer priming and the Retroscript reverse transcription kit (Thermo Fisher Scientific, Cat. #AM1710). Gene expression was quantified by real time RT-PCR, using SYBR Green I master mix (Roche, Cat. #04887352001) and a LightCycler ® 480 device (Roche). mRNA levels were normalized to 18S ribosomal RNA. The primer sets used for all the real time PCR assays are listed in Table S6. In MDA-MB-231, MDA-MB-436, MCF-10A, RMF and HMEC cells, all experiments were repeated three times in triplicate. The data from the three biological replicates were combined, and the mean and standard deviation for each sample was calculated. Statistical comparisons were performed, using GraphPad Prism 8 and the unpaired one tail t-test. In BHMECs the experiments were carried out twice in triplicate, with similar results, and the data from one of the two representative experiments are presented.

**Clinical Proteomic Tumor Analysis Consortium (CPTAC) Data Analysis**

The CPTAC-analyzed breast cancer cohort includes information on 105 TCGA-analyzed samples, of which 25 were derived from patients with basal-like breast cancer and 29, 33 and 18, from patients with luminal A, luminal B and HER-2-enriched breast cancer subtypes respectively [78]. Therefore, the representation of breast cancer subtypes in the CPTAC set of tumors was balanced. Out of the original 105 samples, 102 were tumor samples, of which 28 samples failed to pass a proteomics-based quality control [78]. This left 74 samples for further analyses. Of these, 42 tumors had GSK3α phosphorylated at Ser21 (non-zero abundance) and these tumors were used for calculation of the Spearman correlations between the abundance of GSK3 phosphorylation at Ser9/21 and the abunndance of non-phosphorylated and phosphorylated PP1 and PP2A subunits. The data were retrieved from the CPTAC Breast Cancer Data Portal (https://cptac-data-portal.georgetown.edu/cptacPublic/) [78], that we accessed through the cBioPortal pipeline (https://github.com/cBioPortal/CPTAC-proteomics-pipeline) [79] as was described in an earlier manuscript [80] .

**Reverse Phase Protein Array (RPPA) Data Analysis**

Data were downloaded from https://portal.gdc.cancer.gov/. Overall, 1014 breast cancer patients (all stages) with Reverse Phase Protein Array (RPPA) for ERBB2, ERBB2_pY1248 and ERBB3 were available. The expression of KDM2B was obtained from RNA-Seq data from the same tumor samples. High KDM2B tumors were those with KDM2B expression 1 standard deviation over the mean (728 tumors), and low KDM2B tumors were those with KDM2B expression 1 standard deviation below the mean (164 tumors). Comparisons were statistically analyzed using GraphPad Prism 8 and the unpaired one-tail student's t-test.

**Immunoprecipitation and Immunoblotting**

Cells were lysed in Triton X-100 lysis buffer (20 mM Tris (pH 7.5), 150 mM NaCl, 1 mM EDTA, 1 mM EGTA, 1% Triton X-100) or RIPA buffer (Thermo Scientific, cat. no 89900) supplemented with protease and phosphatase inhibitor cocktails (Sigma-Aldrich, Cat. # 11697498001 and Cat. # 4906837001 respectively). Lysates were sonicated and cleared by centrifugation at 18,000× g for 10 min at 4°C. The cleared lysates were used either for electrophoresis in SDS-PAGE (TritonX-100 and RIPA lysates) or for immunoprecipitation (TritonX-100 Lysates). In the case of immunoprecipitation, the primary antibody was added at the recommended concentration to 500 μl of clarified lysate (1 mg protein) and the mixture was incubated at 4°C, with gentle rocking overnight. The immunoprecipitates were washed three times, 5 min each time, with lysis buffer, and once, also for 5 min, with 0.5M Lithium Chloride. All washes were carried out at 4°C and the washed samples were electrophoresed in SDS-PAGE (20μg protein per lane). Following electrophoresis, cell lysates or immunoprecipitates of cell lysates were transferred to polyvinylidene difluoride membranes in 25 mM Tris, 192 mM glycine. After blocking with 5% nonfat dry milk in TBS and 0.1% Tween-20, the membranes were probed with primary antibodies at the recommended dilution, at 4°C overnight, with gentle rocking. Membrane-bound primary antibodies were detected by horseradish peroxidase-labeled secondary antibodies

(1:2000), and Pierce ECL Western Blotting Substrate (Thermo Scientific, cat. no 32106). All antibodies used in the experiments in this report are listed in Table S8.

**Intracellular Calcium Levels**

Cells were washed twice with Dulbecco's buffered saline (DPBS) without calcium and magnesium (Corning, Cat. #21031CV) and then loaded with Fluo3-AM (Molecular probes, Cat. #F1241) by incubating the cells in a 4 µM Fluo3-AM solution in DPBS at 37°C for 40 minutes. DPBS containing the fluorophore was then discarded and cells were incubated in fresh DPBS for 20 minutes at 37°C. Subsequently, cells were trypsinized and the intensity of fluorescence induced by Ca2+ binding was measured by flow cytometry using the CyAnTM ADP Analyzer (Beckman Coulter) or BD LSRFortessa™ Cell Analyzer.

**Calpain Activity Assay**

Cells were lysed in an EDTA-free lysis buffer containing 20 mM Tris-HCl pH=7.5 and supplemented with a protease inhibitor cocktail (Sigma-Aldrich, Cat. # 4693159001). Subsequently, lysates were mechanically disrupted by forced passage in and out of a syringe and precleared by centrifugation at 18,000× g for 10 min at 4°C. Calpain activity was measured, using the Calpain-Glo assay (Promega, Cat. # G8501) and following the instructions of the manufacturer.

**Calpain-mediated Cleavage of SLUG-WT, Before and After Phosphorylation by GSK3β**

3xFlag-SLUG-WT and 3xFlag-SLUG-4A proteins were immunoprecipitated from 500-1000 µg of lysates from MDAMB-231 cells transduced with the corresponding SLUG constructs. Immunoprecipitates were resuspended in 60 µl kinase buffer (New England Biolabs, Cat. #B6022S), and 30 µl of the total volume was used as the substrate in a GSK3β in vitro phosphorylation reaction. The reaction was carried out at 37°C for 30 minutes by adding 1 µl (500 Units) of purified GSK3β (New England Biolabs, Cat. #P6040S) and ATP (Cell Signaling Technology, Cat. #9804) at a final concentration of 250 µM. The total reaction volume was brought to 40 µl. Following the phosphorylation, the phosphorylation status of the wild-type and 4A mutant SLUG proteins was confirmed by SDS-PAGE and immunoblotting with an anti-phospho-Serine antibody (Abcam, Cat. #AB-9332).

After phosphorylation, the reaction mix was diluted with calpain cleavage buffer (50 mM Tris-HCl, pH 7.4, 50 mM NaCl, 1 mM EDTA, 1 mM EGTA, 1 mM DTT) to a final volume of 100 µl. Thirty µl of this solution was then incubated with or without calpain 1 (Sigma-Aldrich, Cat. #C6108) at a final concentration of 0.3 units/ml, along with 5 mM CaCl2, for 30 seconds at 30°C. Reactions were terminated by adding Laemmli buffer, followed by boiling at 95°C for 10 minutes. The samples were resolved by SDS-PAGE, and the protein cleavage products were detected by immunoblotting.

For the in vitro calpain cleavage assay, the reaction mix was diluted with calpain cleavage buffer (50 mM Tris-HCl, pH 7.4, 50 mM NaCl, 1 mM EDTA, 1 mM EGTA, 1 mM DTT) to a final volume of 100 µl. Thirty µl of this solution was incubated with or without calpain 1 (0.3 units/ml) and 5 mM CaCl2 at 30°C for 30 minutes. Reactions were terminated by adding Laemmli buffer, followed by boiling at 95°C for 10 minutes. The samples were resolved by SDS-PAGE, and the protein cleavage products were detected by immunoblotting.

**Calpain-mediated Cleavage of SLUG-WT, SLUG-4A and SLUG-4D, in vitro.**

pInd20 constructs expressing HA-SLUG-WT, HA-SLUG-4A, and HA-SLUG-4D were transduced into MDA-MB-231 cells, and expression was induced with 1 µg/ml doxycycline for 48 hours. Cells were lysed in Triton X-100 lysis buffer (20 mM Tris (pH 7.5), 150 mM NaCl, 1 mM EDTA, 1 mM EGTA, 1% Triton X-100), supplemented with protease and phosphatase inhibitors (Sigma-Aldrich, Cat. #11697498001 and Cat. #4906837001). Lysates were cleared by centrifugation at 18,000× g for 10 minutes at 4°C.

SLUG variants were immunoprecipitated from the cleared lysates using anti-HA antibodies and incubated with calpain 1 (Sigma-Aldrich, Cat. #C6108) under conditions optimized for calpain activity (2 mM CaCl2, 50 mM Tris (pH 7.5), 1 mM DTT) at 30°C for 30 minutes. Immunoprecipitates were mock-treated or incubated with calpain as controls. Reaction products were resolved by SDS-PAGE and transferred to PVDF membranes. Membranes were probed with the anti-SLUG antibody.

**Flow Cytometry**

Following trypsinization, 500,000 cells were resuspended in 500 µl of PBS containing 3% FBS and 1mM EDTA. Cells were fixed by adding formaldehyde to a final concentration of 4% for 10 min. at 37°C. Subsequently, cells were pelleted, fixation solution was removed, and cells were permeabilized by adding ice-cold 90% methanol and incubating for 30 min on ice. For immunostaining with the anti-phospho-AKT (S473) antibody, methanol was removed, and cells were washed three times with incubation buffer (PBS with 5% Bovine Serum Albumin). After washing, cells were re-suspended in the primary antibody (diluted 1:20 in 100 µl of incubation buffer) and they were incubated for 1 hour at room temperature in the dark. Subsequently, cells were washed with incubation buffer three times, re-suspended in 500 µl of PBS, and analyzed in an LSR-II flow cytometer (Becton Dickinson). For differentiation marker staining, the procedure was performed as described above without fixation or permeabilization. Antibodies were diluted 1:20 and incubation was carried out for 30 minutes at room temperature in the dark.

**Tyrosine Phosphorylation Array**

Tyrosine phosphorylation profiling of control and shKDM2B transduced MDA-MB-231 cells was carried out, using the Full Moon Tyrosine Phosphorylation ProArray (Full Moon BioSystems, Cat. # PST228). Cell lysates were prepared according to the manufacturer's instructions from cells transduced with the shKDM2B (Millipore-Sigma shRNA TRCN0000238779), or the empty vector. Protein concentrations were determined, using the BCA Protein Assay Kit (Thermo Fisher Scientific, Cat. #23227). Fifty µg of total protein from each sample were labeled with biotin (Antibody Array Assay Kit Cat. #KAS02). The biotin-labeled lysates were incubated for 2 hours with array slides containing 228 site-specific phospho-tyrosine antibodies, each spotted in sextuplicate. Following this, arrays were incubated with Cy3-streptavidin (0.5 mg/mL) (Invitrogen Cat. #S-32355) and shipped to Fullmoon BioSystems. The arrays were scanned and analyzed by Fullmoon BioSystems using their standard protocol. Fluorescence intensities were measured to determine site-specific tyrosine phosphorylation changes and normalized using internal controls on the array slides. Array scanning was performed with microarray scanners capable of detecting Cy3 or equivalent fluorophores (Excitation Max: 550 nm; Emission Max: 650 nm). Raw images in TIFF format were provided to the lab for analysis. Image quantification was completed using Fullmoon's Image Analysis Service, ensuring precise and reliable data processing.

**Global and SLUG mRNA-specific Translation In Control and KDM2B Knockdown MDA-MB-231 Cells.**

Cells grown to 60% confluence in 10 cm dishes were starved of L-methionine, L-cystine, and L-glutamine for 1 hour by culturing them in DMEM lacking all three amino acids (Sigma, Cat. #D0422) and supplemented with 10% dialyzed FBS. After starvation, the media was replaced with DMEM containing L-cystine, L-glutamine, and 100 μM L-homopropargylglycine (L-HPG) (Click Chemistry Tools, Cat. #1067), supplemented with 10% FBS. Cells were harvested at the indicated time points by scraping in 1 mL of lysis buffer containing 0.16% SDS in 1xPBS, supplemented with Halt™ Protease and Phosphatase Inhibitor Cocktail (Thermo Fisher Scientific, Cat. #78440). Protein concentration was determined using the Pierce™ BCA Protein Assay Kit (Thermo Fisher Scientific, Cat. #23225). Two mg of protein from each sample were then loaded into a fresh tube, and 1xPBS was added to a final volume of 2 mL. Newly synthesized, L-HPG-labeled proteins in these samples were detected using click chemistry.

Click-chemistry solutions were prepared as 100x stocks and included: Biotin-Azide (10 mM PEG4 carboxamide-6-Azidohexanyl Biotin, Click Chemistry Tools, Cat. #1265), TBTA (10 mM Tris[(1-benzyl-1H-1,2,3-triazol-4-yl) methylamine, Sigma, Cat. #678937), TCEP (100 mM Tris (2-carboxyethyl) phosphine, Sigma, Cat. #C4706), and 100 mM Copper (II) Sulfate (Sigma, Cat. #C1297). TCEP and Copper (II) Sulfate stocks were made fresh the day of the click-chemistry reaction, while Biotin-Azide and TBTA were stored at -80°C. Twenty μL of each stock solution were added to the 2 mL L-HPG-labeled samples, and the mixtures were incubated by shaking for 2 hours in the dark at room temperature. Following this, 10 mL of pre-chilled acetone (-20°C) was added, and samples were placed at -20°C for protein precipitation overnight. The next day, samples were centrifuged at 4000xg for 30 minutes at 4°C. The acetone was removed, and the precipitates were air-dried and resuspended in 700 μL of 1.2% SDS solution containing Halt™ Protease and Phosphatase Inhibitor Cocktail. The samples were then electrophoresed and electroblotted onto

PVDF membranes. The membranes were stained using Revert™ 700 Total Protein Stain (Licor Biosciences, Cat. #296-11020) and probed with an anti-biotin antibody to detect biotinylated nascent peptides.

Changes in SLUG protein levels over time were assessed by immunoblotting. To determine the effects of the KDM2B knockdown on global mRNA translation, newly synthesized protein levels were quantified relative to the total protein levels. To determine the effects of the KDM2B knockdown on SLUG mRNA-specific translation, SLUG protein levels were quantified relative to the levels of global translation efficiency signal.

### TABLE S1

**Correlations between the abundance of total or phosphorylated PP1 subunits and GSK3 phosphorylation at Ser9/21**

| ***PP1 subunits*** | ***R**** | ***Phosphorylated PP1 subunits*** | ***R**** |
|---|---|---|---|
| PPP1CA | -0.66 | PPP1CA_pT320 | 0.76 |
| | | PPP1CA_pS22 | 0.39 |
| PPP1CB | -0.64 | PPP1CB_pT316 | 0.88 |
| PPP1R7 | -0.51 | PPP1R7_pS12 | 0.83 |
| | | PPP1R7_pS322 | 0.63 |
| | | PPP1R7_pS27 | 0.69 |

* ***R*** represents the Spearman correlation coefficient between the indicated PP1 subunits and GSK3 phosphorylated at Ser9/21. Note the negative correlations of the unmodified subunits and the positive correlations of the phosphorylated subunits. These data suggest that the indicated unmodified subunits dephosphorylate GSK3 phosphorylated at Ser9/21 and that phosphorylation of the indicated subunits renders PP1 inactive against GSK3 phosphorylated at this site.

**TABLE S2**
**Correlations between the abundance of total or phosphorylated PP2 subunits and GSK3 phosphorylation at Ser9/21**

| ***PP2A Subunits*** | ***R**** | ***Phosphorylated PP2A subunits*** | ***R**** |
|---|---|---|---|
| PPP2CA | -0.53 | | NC |
| PPP2CB | -0.54 | | NC |
| PPP2R1A | -0.4 | | NC |
| PPP2R1B | -0.55 | | NC |
| PPP2R2D | -0.56 | | NC |
| PPP2R5D | ?? | PPP2R5D_pS573 | 0.74 |
| PPPR5E | NC | PPP2R5E_pS32 | 0.78 |
| PPPR5A | NC | PPP2R5A_pS49 | 0.74 |

***** ***R*** represents the Spearman correlation coefficient between the indicated PP2A subunits and GSK3 phosphorylated at Ser9/21. Note the negative correlations of the unmodified subunits and the positive correlations of the phosphorylated subunits. These data suggest that the indicated unmodified subunits dephosphorylate GSK3 phosphorylated at Ser9/21 and that phosphorylation of the indicated subunits renders PP2A inactive against GSK3 phosphorylated at this site. (NC stands for No Correlation).

**Table S3: Cell lines and corresponding growth media.**

| Cell line | Growth Medium |
|---|---|
| **HEK293T** | DMEM high glucose, 10% FBS, P/S, non-essential amino acids, L-glutamine |
| **MDAMB-231** | DMEM high glucose, 10% FBS, P/S, non-essential amino acids, L-glutamine |
| **MCF-10A** | MEGM Mammary Epithelial Cell Growth Medium, Lonza, cat no cc-3150 |
| **MDAMB-436** | DMEM high glucose, 10% FBS, P/S, non-essential amino acids, L-glutamine |
| **621-HMEC** | MEGM Mammary Epithelial Cell Growth Medium, Lonza, cat no cc-3150 |
| **629- BHMEC** | MEGM Mammary Epithelial Cell Growth Medium, Lonza, cat no cc-3150 |
| **RMF-WT** | DMEM high glucose, 10% FBS, P/S, non-essential amino acids, L-glutamine |

**Table S4: Inhibitors, growth factors and chemicals used for cell culture**

| Compound | Brand | Cat. # | Use |
|---|---|---|---|
| **AZD5363** | Selleckchem | S8019 | pan AKT inhibitor |
| **RO 31-8220** | Tocris | 2002 | Classical PKC inhibitor |
| **GSK 650394** | Tocris | 3572 | pan SGK inhibitor |
| **PF 4708671** | Tocris | 4032 | p70S6K inhibitor |
| **BI-D1870** | Enzo life sciences | BML-EI407-0001 | p90RSK inhibitor |
| **AR-A 014418** | Tocris | 3966 | GSK3 inhibition |
| **Daminozide** | Tocris | 4684 | KDM2B demethylase inhibition |
| **Chloroquine** | Sigma-Aldrich | C6628 | Autophagy inhibitor |
| **3-Methyladenine** | Sigma-Aldrich | M9281 | Autophagy inhibitor |
| **Concanamycin A** | Sigma-Aldrich | C9705 | Autophagy inhibitor |
| **Cycloheximide** | Sigma-Aldrich | C4859 | Protein synthesis inhibitor |
| **Bortezomib** | Selleckchem | S1013 | Proteasome inhibitor |
| **Ionomycin** | Cell Signaling Technology | 9995 | Calpain activation |
| **Human TGF-β1** | Cell Signaling Technology | 8915 | EMT induction |
| **Acetyl-Calpastatin (184-210)** | Tocris | 2950 | Calpain inhibition |
| **Osimertinib** | MedChemExpress | HY-15772 | EGFR inhibitor |
| **Erlotinib** | MedChemExpress | HY-50896 | EGFR tyrosine kinase inhibitor |
| **Doxycycline** | MedChemExpress | HY-N0565 | Induction of SLUG-WT, SLUG-4A, or SLUG-4D expression in Pin20-transduced cells |

**Table S5: Cloning and mutagenesis primers**

| Primer | Use | Sequence 5´→3´ |
|---|---|---|
| **FLAG-SLUG** | Subcloning | F: GGG GAA TTC A ATG CCG CGC TCC TTC CTG GTC<br>R: GGG GAA TTC TCA CTA CTT GTC ATC GTC ATC |
| **SLUG-4A/4D Gene Block** | Assembly/Mutagenesis | F: TCC TCA GCT CAG GAG CAT ACA GCC CCATC<br>R: TGC ATA AAT TGC ACT GAA ACT TTT CAG CTT CAA TG |
| **pENTR-SLUG** | Assembly/Mutagenesis | F: GTT TCA GTG CAA TTT ATG CAA TAA GAC CTA TTC AAC TTT TTC<br>R: GTA TGC TCC TGA GCT GAG GAT CTC TGG TTG |
| **GSK3β-Myc** | Cloning | F: GCG CTA CGT AAT GTC AGG GCG GCC CAG AAC C<br>R: GCG CGA ATT CTC ACA GGT CCT CCT CGC TGA TCA GCT TCT GCT CGG TGG AGT TGG AAG CTG |
| **GSK3β-Myc** | Subcloning | F: CAC CTC AGG GCG GCC C<br>R: CAG GTC CTC CTC GC |
| **GSK3β-S9A** | Mutagenesis | F: CTC TCC GCA AAG GCG GTG GTT CTG GGC<br>R: GCC CAG AAC CAC CGC CTT TGC GGA GAG |

### Table S6: Real-Time RT-PCR primers

| Primer | Sequence 5´→3´ |
|---|---|
| **Ribosomal 18S** | F: GTA ACC CGT TGA ACC CCA TT<br>R: CCA TCC AAT CGG TAG TAG CG |
| **KDM2B** | F: TCT ACG AGA TCG AGG ACA GGA<br>R: ACC AGC ACA TCT CAT AGT AGA AGG |
| **SLUG** | F: TGG TTG CTTCAAGGACACAT<br>R: CAG AAT GGG TCT GCA GAT GA |
| **SOX9** | F: GCT CTG GAG ACT TCT GAA CGA<br>R:CCG TTC TTC ACC GAC TTC CT |
| **CALPASTATIN** | F: TCC AGA ACC TAT GCT GGT GG<br>R: TTA CAG CCT TGG TTT CTG TGG |
| **SNAIL** | F: CCA GTG CCT CGA CCA CTA TG<br>R: CTG CTG GAA GGT AAA CTC TGG A |

### Table S7: Cloning and mutagenesis gene blocks

| Gene Block | Sequence 5´→3´ |
|---|---|
| SLUG-4A | AGGAGCATACAGCCCCATCACTGTGTGGACTACCGCTGCTCCATTCCACGCCCAGCTACCC<br>AATGGCCTCTCTCCTCTTTCCGGATACTCCTCATCTTTGGGGCGAGTGAGTCCCCCTCCTC<br>CAGCCGACACCTCCGCCAAGGACCACGCCGGCTCAGAAGCCCCCATTAGTGATGAAGAGGA<br>AAGACTACAGTCCAAGCTTTCAGACCCCCATGCCATTGAAGCTGAAAAGTTTCAGTGC |
| SLUG-4D | AGGAGCATACAGCCCCATCACTGTGTGGACTACCGCTGCTCCATTCCACGCCCAGCTACCC<br>AATGGCCTCTCTCCTCTTTCCGGATACTCCTCATCTTTGGGGCGAGTGAGTCCCCCTCCTC<br>CAGATGACACCTCCGACAAGGACCACGATGGCTCAGAAGACCCCATTAGTGATGAAGAGGA<br>AAGACTACAGTCCAAGCTTTCAGACCCCCATGCCATTGAAGCTGAAAAGTTTCAGTGC |

### Table S8: Antibodies

| Antibody | Use | Brand | Cat. # |
|---|---|---|---|
| **KDM2B** | WB | Millipore | 09-864 |
| **SLUG** | WB | Cell Signaling Technology | 9585 |
| **SOX9** | WB | Millipore | AB5535 |
| **SNAIL** | WB | Cell Signaling Technology | 3879 |
| **Phospho-GSK-3α/β (Ser21/9)** | WB | Cell Signaling Technology | 8566 |
| **GSK-3α/β** | WB | Cell Signaling Technology | 5676 |
| **Phospho-GSK-3α (Ser21)** | WB | Cell Signaling Technology | 9316 |
| **GSK-3α** | WB | Cell Signaling Technology | 9316 |
| **Phospho-GSK-3β** | WB | Cell Signaling Technology | 5558 |
| **GSK-3β** | WB | Cell Signaling Technology | 12456 |
| **Calpain 1** | WB | Cell Signaling Technology | 2556 |
| **Calpain 2** | WB | Cell Signaling Technology | 2539 |
| **Calpain 9** | WB | Santa Cruz Biotechnology | sc-166750 |
| **Calpastatin** | WB | Cell Signaling Technology | 4146 |
| **INPP4B** | WB | Cell Signaling Technology | 1454 |
| **Phospho AKT (S473)** | WB | Cell Signaling Technology | 4060 |
| **Phospho AKT (T308)** | WB | Cell Signaling Technology | 2965 |
| **Phospho Tyrosine** | WB | Millipore | 05-321 |
| **Phospho Serine** | WB | Abcam | AB-9332 |
| **PKC phospho substrate** | WB | Cell Signaling Technology | 2261 |
| **E-cadherin** | WB | Cell Signaling Technology | 3195 |
| **Vimentin** | WB | Cell Signaling Technology | 5741 |
| **GAPDH** | WB | Millipore | MAB374 |
| **α-Tubulin** | WB | Sigma-Aldrich | T5168 |

| | | | |
|---|---|---|---|
| **Phospho-Akt (Ser473) - Alexa Fluor® 488** | FC | Cell Signaling Technology | 4071 |
| **SGK1** | WB | Cell Signaling Technology | 12103 |
| **SGK2** | WB | Cell Signaling Technology | 5595 |
| **Phospho-SGK3 (Thr320)** | WB | Cell Signaling Technology | 5642 |
| **SGK3** | WB | Cell Signaling Technology | 8156 |
| **HA-Tag** | WB | Cell Signaling Technology | 3724 |
| **Vinculin** | WB | Cell Signaling Technology | 4650 |
| **STAT4** | WB | Cell Signaling Technology | 2653 |
| **HER4/ErbB4** | WB | Cell Signaling Technology | 4795 |
| **PTK2B(Pyk2)** | WB | Cell Signaling Technology | 43550 |
| **FGF Receptor 1** | WB | Cell Signaling Technology | 9740 |
| **Biotin-HRP** | WB | Cell Signaling Technology | 7075 |
| **TEK(Tie2)** | WB | Cell Signaling Technology | 7403 |
| **α-Actinin** | WB | Cell Signaling Technology | 3134 |

### Table S9: shRNA Constructs

| Target Gene | Brand | Cat. # |
|---|---|---|
| **KDM2B** | Dharmacon | TRCN0000118437 |
| **KDM2B** | Millipore-Sigma | TRCN0000238779 |
| **STAT4** | Millipore-Sigma | TRCN0000020894 |
| **STAT4** | Millipore-Sigma | TRCN0000020895 |
| **FGFR1** | Millipore-Sigma | TRCN0000312574 |
| **TEK** | Millipore-Sigma | TRCN0000000412 |
| **PTK2B** | Millipore-Sigma | TRCN0000231523 |
| **Erbb4/HER4** | Millipore-Sigma | TRCN0000001411 |
| **MERTK** | Millipore-Sigma | TRCN0000000865 |
| **Calpastatin** | Dharmacon | TRCN0000073638 |

### SUPPLEMENTARY FIGURE LEGENDS

#### Figure S1: Posttranscriptional Regulation and Protein Destabilization of SLUG by KDM2B

**A.** Quantitative RT-PCR failed to detect consistent downregulation of SLUG, SNAIL and SOX9 in shKDM2B-transduced MDA-MB-436, HMEC, BHMEC and RMF cells. **B.** Quantification of the western blots in figure 2A confirmed that the degradation of SLUG is more rapid in shKDM2B-transduced, than in shControl transduced MDA-MB-231 (Left panel) and MCF-10A cells (Right panel). Data presented as Mean±SD of values from three independent experiments. Asterisks indicate p-value ≤0.05. **C.** The translational efficiency of the SLUG mRNA is not significantly altered by the knockdown of KDM2B. Coupled RNA-seq and ribosome profiling data in control and KDM2B knockdown MDA-MB-231 cells, allowed us to measure the shKDM2B-induced change in translational efficiency (ΔTE) of different mRNAs genome wide. Table shows the ΔTE of the SLUG mRNA, which was not significantly affected by the KDM2B knockdown and the ΔTE and four additional mRNAs whose translational efficiency was significantly reduced. **D.** Control and shKDM2B-transduced MDA-MB-231 cells were labeled with the methionine analog L-HPG, after one hour of methionine starvation. *(Left upper panel)* Newly synthesized proteins were detected by probing western blots of cell lysates, harvested at the indicated time points, with biotin-azide, which reacts with the alkyne group of L-HPG in labeled proteins via click chemistry. *(Left middle panel).* Total SLUG protein levels in control and shKDM2B cells were measured by probing a western blot of the same lysates with an anti-SLUG antibody. *(Left lower panel).* Total protein levels in the same lysates were visualized by staining with the Revert Protein Stain. *(Right upper panel)*. The knockdown of KDM2B induces global translational repression. The L-HPG signal, which measures the abundance of newly synthesized proteins (upper left panel), was normalized to the total protein signal (left lower panel) for in each time point and condition. Simple linear regression with line of best fit calculation was performed to determine the relative rates of protein production in arbitrary units. Line of best fit was plotted with the 95% confidence interval displayed as a red or blue backdrop in the two data series. *(Right lower panel)* The SLUG western blot signal (left middle panel) was normalized for each time point and condition to the global translational efficiency signal in the upper right panel. Simple linear regression with calculation of the line of best fit was performed to determine the relative rates of Slug/Global Translational Efficiency in arbitrary units.

**Figure S2: GSK3β inactivation promotes the differentiation of MCF10A and MDA-MB-231 cells in culture and inhibits TGFβ-induced EMT in MCF10A cells.**

**A.** Probing immunoblots of cell lysates derived from MDA-MB-231 and MCF-10A cells transduced with shControl, or shKDM2B lentiviral constructs with anti-phospho-GSK3α (Ser21) (Left panel), or anti-phospho-GSK3β (Ser9) (Right panel) antibodies, revealed that shKDM2B promotes the phosphorylation of both, although the phosphorylation of GSK3β is more robust. Antibodies to total GSK3α, GSK3β and tubulin were used as controls. **B.** *(Left panel)* MCF10A cells, which are known to express basal/myoepithelial markers [79], were treated with the GSK3 inhibitor AR-A-14418 (2 μM) for seven days and they were harvested at the indicated time points. Probing the cell lysates with the indicated antibodies, revealed a robust, but transient downregulation of SLUG at day 2, a gradual upregulation of E-Cadherin and a gradual downregulation of vimentin, suggesting that GSK3 inhibition induces mesenchymal to epithelial transition (MET). *(Middle panel).* MCF-10A cells were treated with TGFβ (4 ng/ml) and they were harvested at the indicated time points. Probing immunoblots of the lysates with the indicated antibodies, confirmed that TGFβ induces EMT, as determined by the gradual upregulation of SLUG and Vimentin and the gradual downregulation of E-Cadherin. *(Right panel)* Repeating the experiment in the middle panel in the presence of AR-A 14418, revealed that inhibition of GSK3 interferes with the induction of EMT by TGFβ. **C.** MDAMB-231 cells were treated with AR-A-14418 (2 μM) or DMSO. Seven days later the cells were stained with the indicated FITC- and PE or APC-conjugated antibodies (CD44/CD24, CD49f/EpCAM or CD49f/CD24) and they were analyzed by flow-cytometry.

**Figure S3: KDM2B Modulates GSK3β Phosphorylation Through Kinase Activation and Dephosphorylation Inhibition**

**A.** MDA-MB-231 and MCF-10A cells transduced with shControl or shKDM2B constructs, were treated with the indicated kinase inhibitors (pan AKT, AZD-5363 (0.5 μM); Classical PKC, RO-31-8220 (0.1 μM); p70S6K, PF-4708671 (10 μM) and p90RSK, BI-D1870 (10 μM). Probing immunoblots of cell lysates harvested at two hours from the start of the treatment, revealed that AKT and PKC phosphorylate GSK3 in both the shControl and shKDM2B cells. **B.** Probing with the same antibodies cell lysates of the same cells, following a 24-hour treatment with the SGK inhibitor GSK650394 (10 μM), revealed that SGK also promotes the steady state phosphorylation of GSK3. **C.** The six cell lines in our mammary cell line panel were transduced with shControl or shKDM2B lentiviral constructs. Probing immunoblots of cell lysates of these cell lines with anti-phospho-AKT (Thr308 and Ser473) and anti-Total AKT antibodies, revealed that shKDM2B promotes AKT activation in some cell lines (MDA-MB-231 and MCF-10A) and deactivation in others (MDA-MB-436, HMEC, BHMEC and RMF). **D.** Probing immunoblots of the same lysates with anti-INPP4B and anti-GAPDH (loading control) antibodies, revealed that the expression of INPP4B exhibits a perfect negative correlation with the phosphorylation of AKT. **E.** Probing immunoblots of the same lysates with a classical PKC phospho-substrate antibody, revealed that PKC is inhibited in shKDM2B-transduced cells. **F.** Probing immunoblots of cell lysates from five of the cell lines in our cell line panel, with anti-SGK1, anti-SGK2, anti-SGK3, and anti-phospho-SGK3 antibodies, shows that shKDM2B promotes the upregulation of SGK2 in in three cell lines and the phosphorylation of SGK3 in two. **G.** MDA-MB-231 cells were treated with the pan-AKT inhibitor AZD-5363 (5 μM) and they were lysed and harvested at the indicated time points. Probing immunoblots of these lysates with anti-phospho-GSK3β and anti-Total GSK3β antibodies, provided evidence that shKDM2B stabilizes GSK3β phosphorylation.

**Figure S4: The shKDM2B-induced degradation of SLUG is calpain-mediated.**

**A.** MDA-MB-231 cells transduced with shControl or shKDM2B constructs were treated with chloroquine (CQ, 30 μM), 3-methyladenine (3-MA, 10 mM), or concanamycin A (1 μM) for 24 hours. Probing immunoblots of cell lysates harvested at the indicated time points from the start of the treatment, with anti-SLUG or anti-Tubulin (loading control) antibodies, revealed that none of the autophagy inhibitors rescued the shKDM2B-induced destabilization of SLUG. **B.** All six cell lines in our panel were transduced with shControl or shKDM2B. Probing immunoblots of lysates derived from these cells with anti-calpain 1, 2 or 9 antibodies, revealed that whereas Calpain 1 and 2 were upregulated in some cell lines, calpain 9 was downregulated in all. **C.** Measurement of the calpastatin mRNA levels by quantitative RT-PCR in shControl and shKDM2B-transduced MDA-MB-436, HMEC, BHMEC and RMF cells, confirmed that calpastatin is downregulated by KDM2B at the RNA level in all the cell lines, except of HMEC. Data are presented as mean ± SD and asterisks indicate statistical significance (p-value ≤ 0.05). **D.** MDAMB-231 and MCF-10A cells were treated with increasing concentrations of ionomycin, as indicated. Calpain 1 and 2 activity was measured in cell lysates harvested at 1 hour from the start of the ionomycin treatment, using a luminescent assay. Data are presented as mean ± SD and asterisks indicate statistical significance (* p-value ≤ 0.05, ** p-value ≤ 0.01, *** p-value ≤ 0.001).

**Figure S5: The RTKs TEK and MERTK, and the transcription factor STAT4, do not contribute to the shKDM2B-induced GSK3 phosphorylation and SLUG degradation phenotype.**

**A.** Tyrosine phosphorylation profiling of control and KDM2B knockdown MDA-MB-231 cells. Cell lysates were used to probe the Full Moon Tyrosine Phosphorylation ProArray. All antibodies in the array were included in sextuplicate. Images were quantified and the change in the intensity of the images in the KDM2B knockdown relative to the control cells was calculated. **B.** The phosphorylation of the RTKs TEK and MERTK does not contribute to the shKDM2B-induced GSK3 phosphorylation and SLUG degradation phenotype. Control and KDM2B knockdown MDA-MB-231 cells, were transduced with shTEK, and shMERTK constructs, or with the shControl, and their lysates were probed with the indicated antibodies. **C.** The phosphorylation of the transcription factor STAT4 does not contribute to the shKDM2B-induced GSK3 phosphorylation and SLUG degradation phenotype. Control and KDM2B knockdown MDA-MB-231 cells, were transduced with an shSTAT4, construct, or with the shControl, and their lysates were probed with the indicated antibodies.

**Figure S6: Upregulation of EGFR Ligands and ERBB Family Proteins in the Context of KDM2B Knockdown**

**A.** Heatmap of cytokine and growth factor mRNAs in control and KDM2B knockdown MDA-MB-231 cells. The heatmap was based on the results of an RNA-seq study of control and shKDM2B-transduced MDA-MB-231 cells. Several ligands, including HB-EGF, EREG, BTC, EGF, TGF-α, and AREG, were upregulated in shKDM2B cells (see text for details) **B.** Distribution of the abundance of ERBB2, ERBB2_pY1248 and ERBB3 in human mammary adenocarcinomas expressing high (Red) or low (Blue) levels of KDM2B. Horizontal lines indicate the mean values. Tumors with low KDM2B tend to express higher levels of ERBB2, ERBB2_pY1248 and ERBB3, than tumors with high KDM2B. Statistical analysis was performed using the unpaired one tail student's t test.

**Figure S7: The knockdown of KDM2B in human mammary gland-derived cell lines activates GSK3β phosphorylation and calpain activation pathways via paracrine mechanisms.**

Model of the calpain-dependent degradation of SLUG induced by the knockdown of KDM2B. The knockdown of KDM2B results in the indirect activation of tyrosine kinase receptor-dependent paracrine mechanisms. The tyrosine kinase receptor-initiated signals activate AKT and/or SGK, which phosphorylate GSK3 at Ser9/21. In addition, they block the activities of PP1 and PP2A toward GSK3, phosphorylated at Ser9/21. The ultimate output of these two pathways converging on GSK3 is the enhanced phosphorylation of GSK3 at Ser9/21. The tyrosine kinase receptor signals also induce Ca2+ influx and calpastatin (CAST) downregulation and the combination of these two events results in the activation of calpains 1 and 2. Under steady state conditions, SLUG is phosphorylated by GSK3, and the phosphorylated SLUG undergoes degradation via the proteasome. Inactivation of GSK3 by phosphorylation at Ser9/21 induced by the knockdown of KDM2B, prevents the slow proteasomal degradation of SLUG, but renders SLUG sensitive to degradation by calpains 1 and 2. This results in a switch from the slow proteasomal degradation to the rapid calpain-dependent degradation.

A

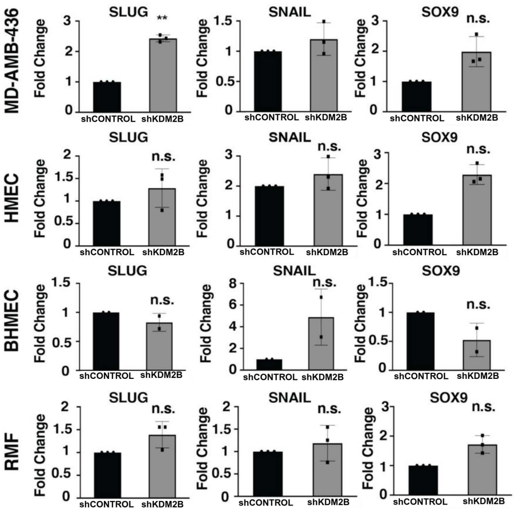


B

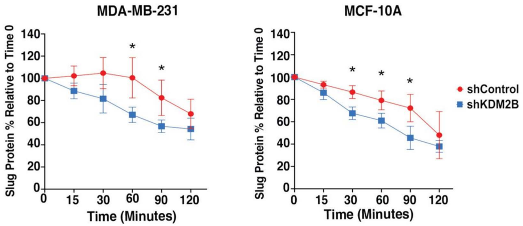


C

| Gene | Log2FC ΔTE | Adjusted P-Value |
|---|---|---|
| SLUG | -0,28 | 0,21 |
| CTNNB1 | -0,58 | 0,004 |
| PNP | -0,70 | 0,004 |
| PSMC2 | -0,59 | 0,004 |

D

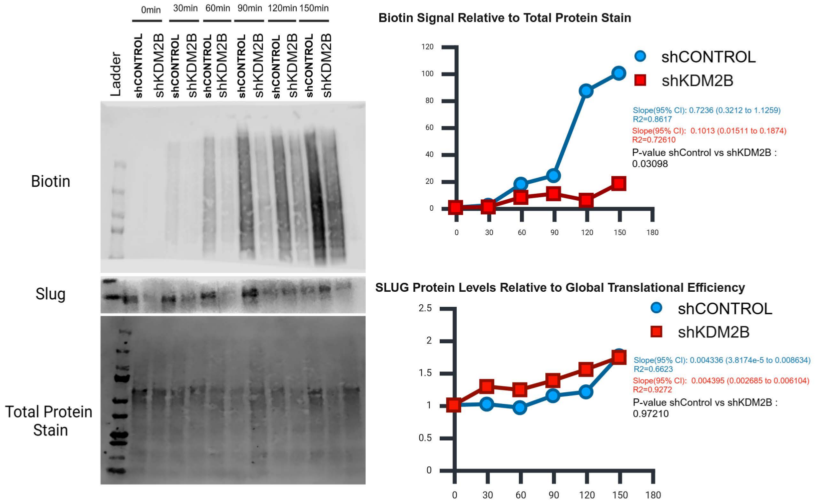

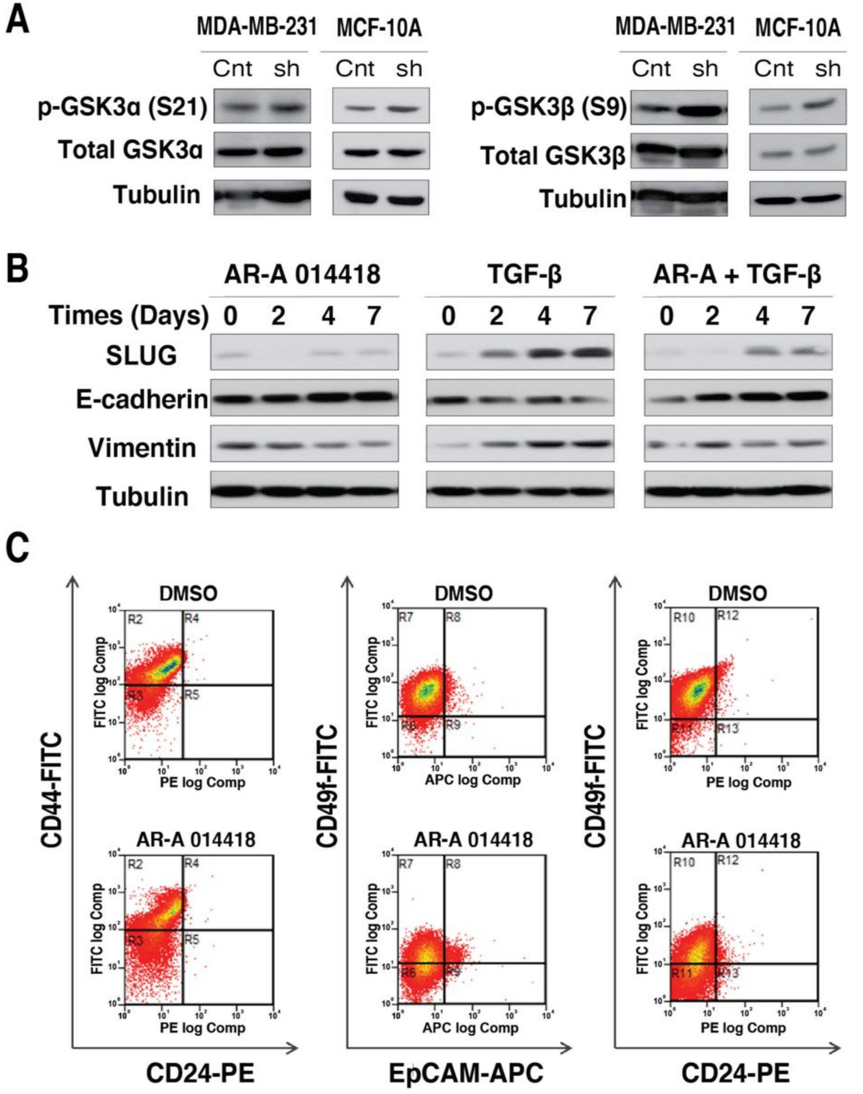
A
MDA-MB-231
MCF-10A
Cnt sh
p-GSK3α (S21)
Total GSK3α
Tubulin
p-GSK3β (S9)
Total GSK3β
B
AR-A 014418
TGF-β
AR-A + TGF-β
Times (Days) 0 2 4 7
SLUG
E-cadherin
Vimentin
Tubulin
C
DMSO
AR-A 014418
CD44-FITC
CD24-PE
CD49f-FITC
EpCAM-APC
FITC log Comp
PE log Comp
APC log Comp
R2 R3 R4 R5
R7 R8 R6 R9
R10 R11 R12 R13

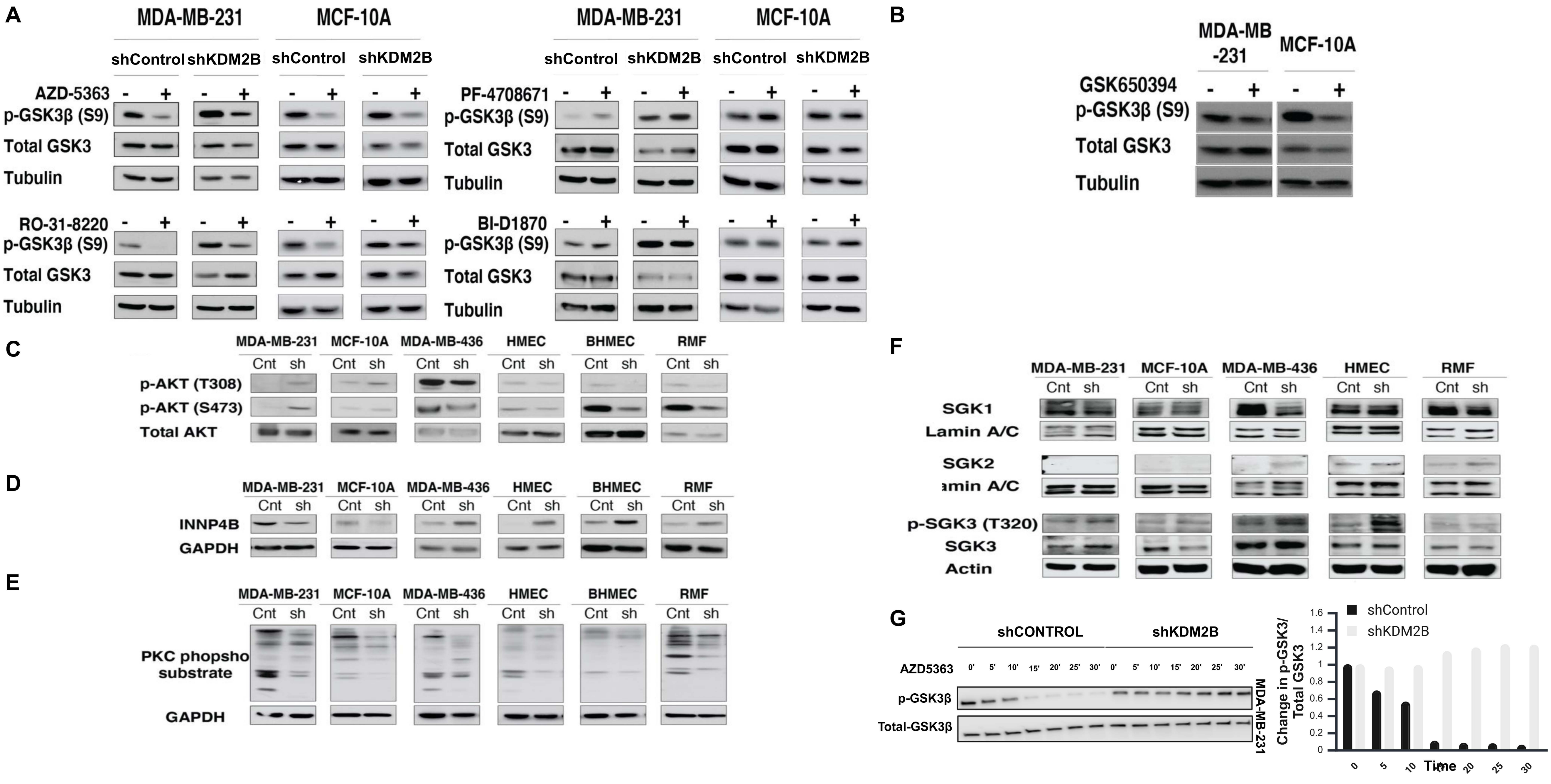
A
MDA-MB-231
MCF-10A
shControl
shKDM2B
AZD-5363
RO-31-8220
PF-4708671
BI-D1870
p-GSK3β (S9)
Total GSK3
Tubulin
B
MDA-MB -231
MCF-10A
GSK650394
p-GSK3β (S9)
Total GSK3
Tubulin
C
MDA-MB-231
MCF-10A
MDA-MB-436
HMEC
BHMEC
RMF
Cnt
sh
p-AKT (T308)
p-AKT (S473)
Total AKT
D
INNP4B
GAPDH
E
PKC phopsho substrate
GAPDH
F
SGK1
Lamin A/C
SGK2
amin A/C
p-SGK3 (T320)
SGK3
Actin
G
shCONTROL
shKDM2B
AZD5363
0'
5'
10'
15'
20'
25'
30'
p-GSK3β
Total-GSK3β
MDA-MB-231
shControl
shKDM2B
Change in p-GSK3/ Total GSK3
Time
1.6
1.4
1.2
1
0.8
0.6
0.4
0.2
0
0
5
10
20
25
30

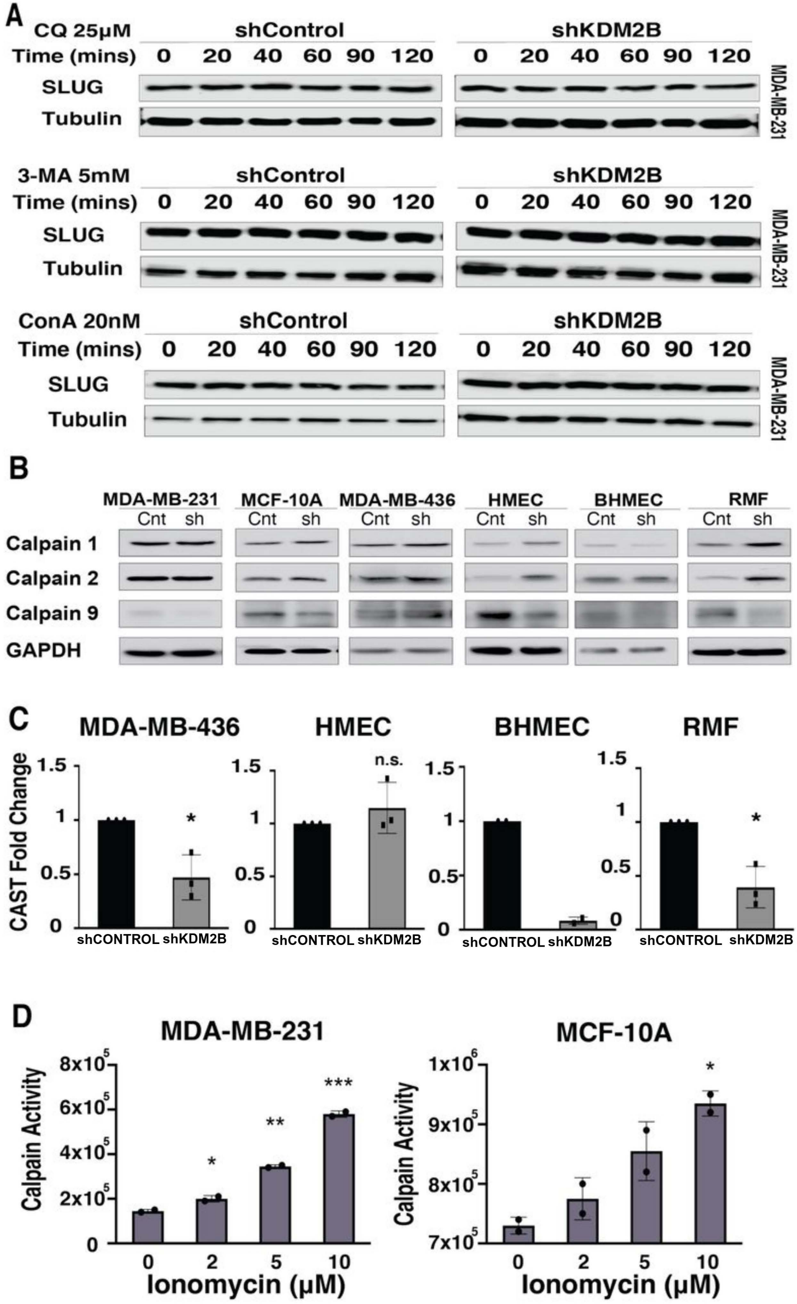
A
CQ 25μM
shControl
shKDM2B
Time (mins) 0 20 40 60 90 120 0 20 40 60 90 120
SLUG
Tubulin
MDA-MB-231
3-MA 5mM
shControl
shKDM2B
Time (mins) 0 20 40 60 90 120 0 20 40 60 90 120
SLUG
Tubulin
MDA-MB-231
ConA 20nM
shControl
shKDM2B
Time (mins) 0 20 40 60 90 120 0 20 40 60 90 120
SLUG
Tubulin
MDA-MB-231
B
MDA-MB-231 MCF-10A MDA-MB-436 HMEC BHMEC RMF
Cnt sh Cnt sh Cnt sh Cnt sh Cnt sh Cnt sh
Calpain 1
Calpain 2
Calpain 9
GAPDH
C
MDA-MB-436
HMEC
BHMEC
RMF
CAST Fold Change
1.5
1
0.5
0
*
n.s.
shCONTROL shKDM2B
D
MDA-MB-231
MCF-10A
Calpain Activity
8x10^5
6x10^5
4x10^5
2x10^5
0
1x10^6
9x10^5
8x10^5
7x10^5
*
**
***
0 2 5 10
Ionomycin (μM)

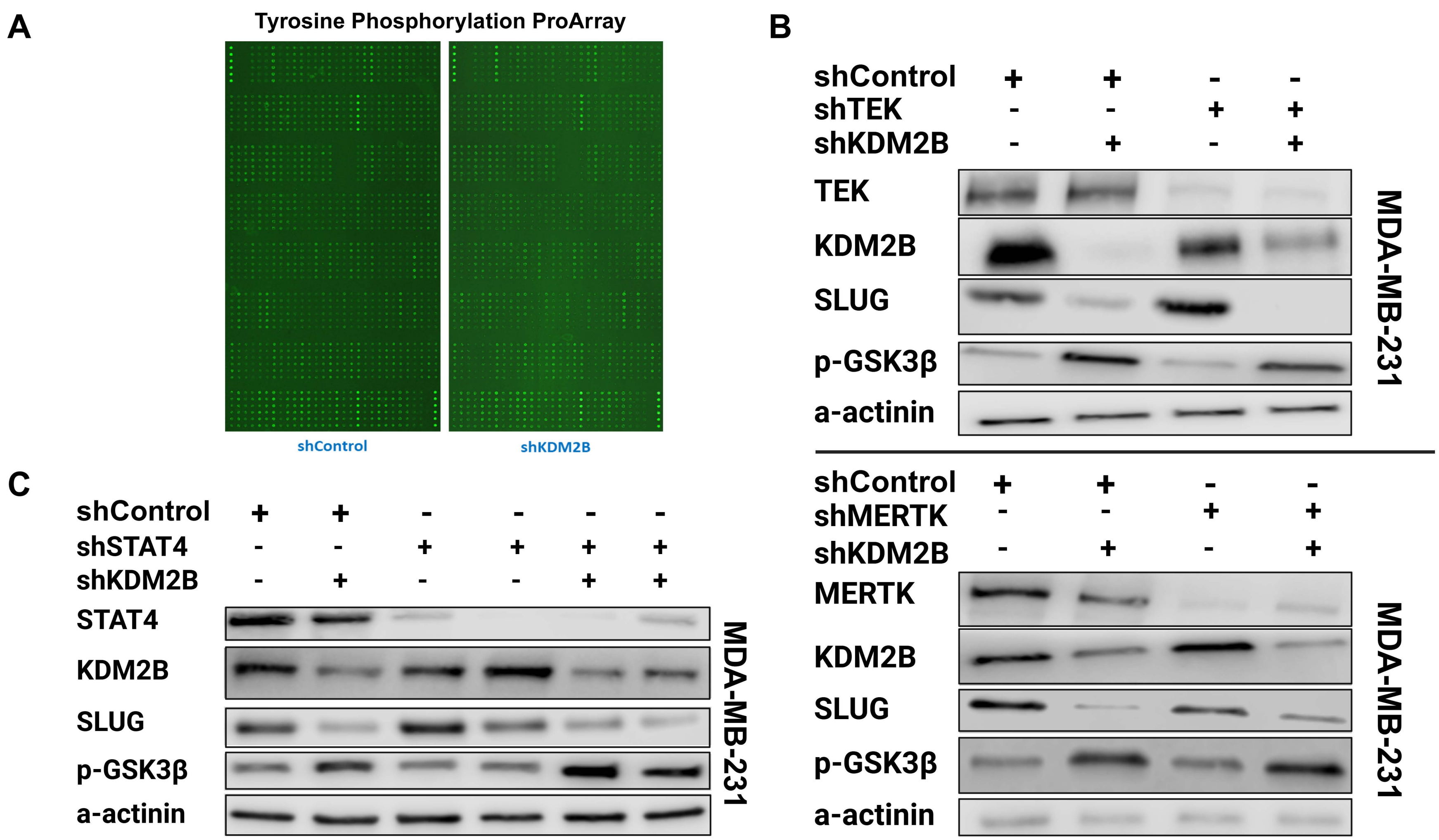

A
Tyrosine Phosphorylation ProArray
shControl
shKDM2B
B
shControl + + - -
shTEK - - + +
shKDM2B - + - +
TEK
KDM2B
SLUG
p-GSK3β
a-actinin
MDA-MB-231
shControl + + - -
shMERTK - - + +
shKDM2B - + - +
MERTK
KDM2B
SLUG
p-GSK3β
a-actinin
MDA-MB-231
C
shControl + + - - - -
shSTAT4 - - + + + +
shKDM2B - + - - + +
STAT4
KDM2B
SLUG
p-GSK3β
a-actinin
MDA-MB-231

A

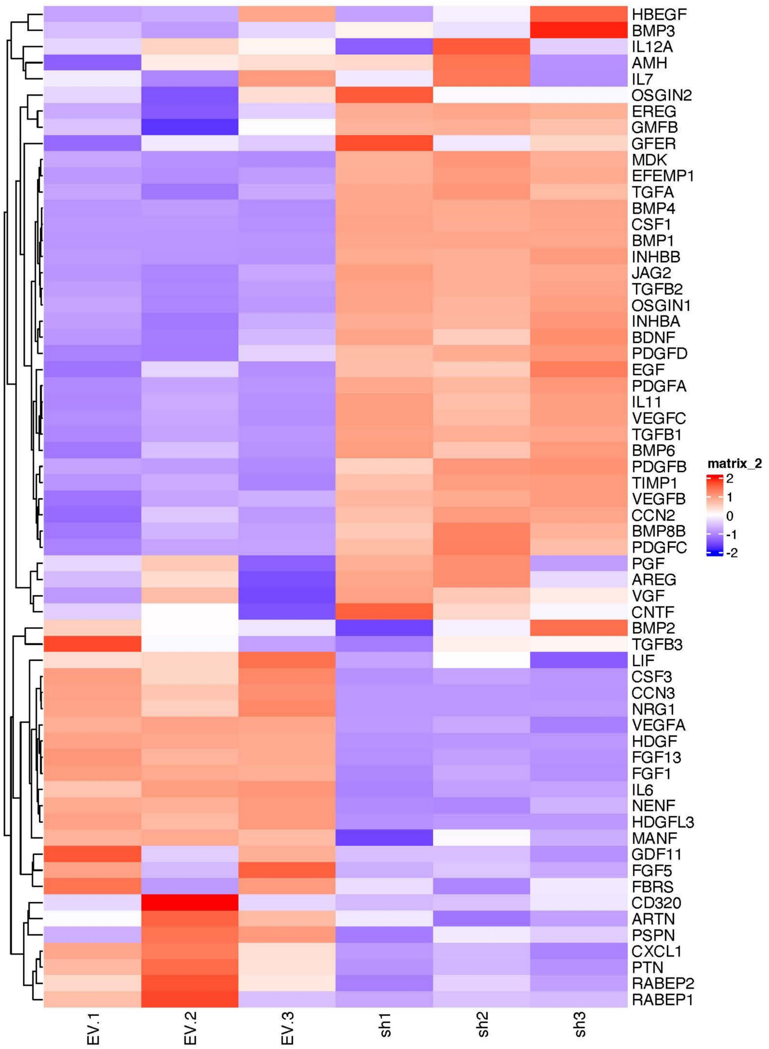

HBEGF
BMP3
IL12A
AMH
IL7
OSGIN2
EREG
GMFB
GFER
MDK
EFEMP1
TGFA
BMP4
CSF1
BMP1
INHBB
JAG2
TGFB2
OSGIN1
INHBA
BDNF
PDGFD
EGF
PDGFA
IL11
VEGFC
TGFB1
BMP6
PDGFB
TIMP1
VEGFB
CCN2
BMP8B
PDGFC
PGF
AREG
VGF
CNTF
BMP2
TGFB3
LIF
CSF3
CCN3
NRG1
VEGFA
HDGF
FGF13
FGF1
IL6
NENF
HDGFL3
MANF
GDF11
FGF5
FBRS
CD320
ARTN
PSPN
CXCL1
PTN
RABEP2
RABEP1
matrix_2
2
1
0
-1
-2
EV.1
EV.2
EV.3
sh1
sh2
sh3


B

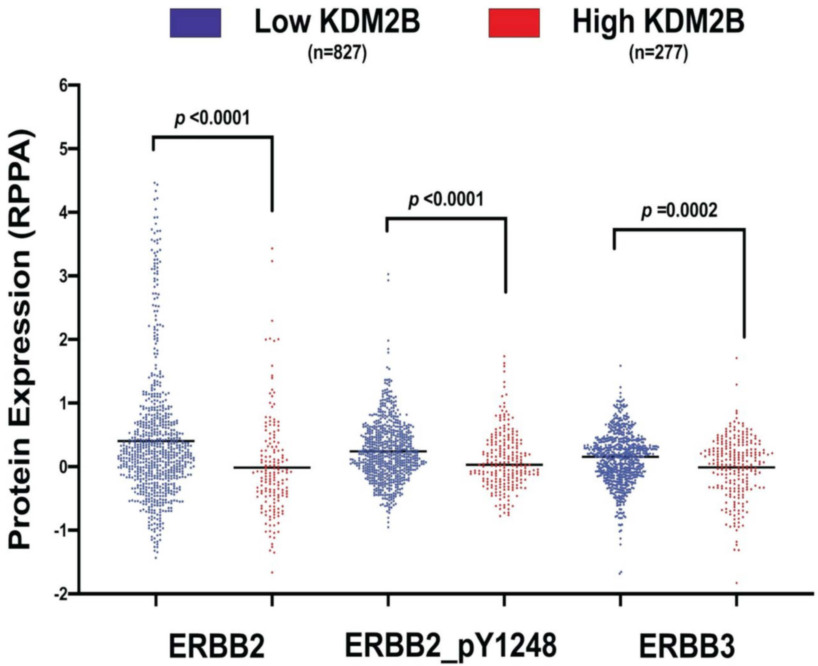

Low KDM2B
(n=827)
High KDM2B
(n=277)
Protein Expression (RPPA)
6
5
4
3
2
1
0
-1
-2
p <0.0001
p <0.0001
p =0.0002
ERBB2
ERBB2_pY1248
ERBB3

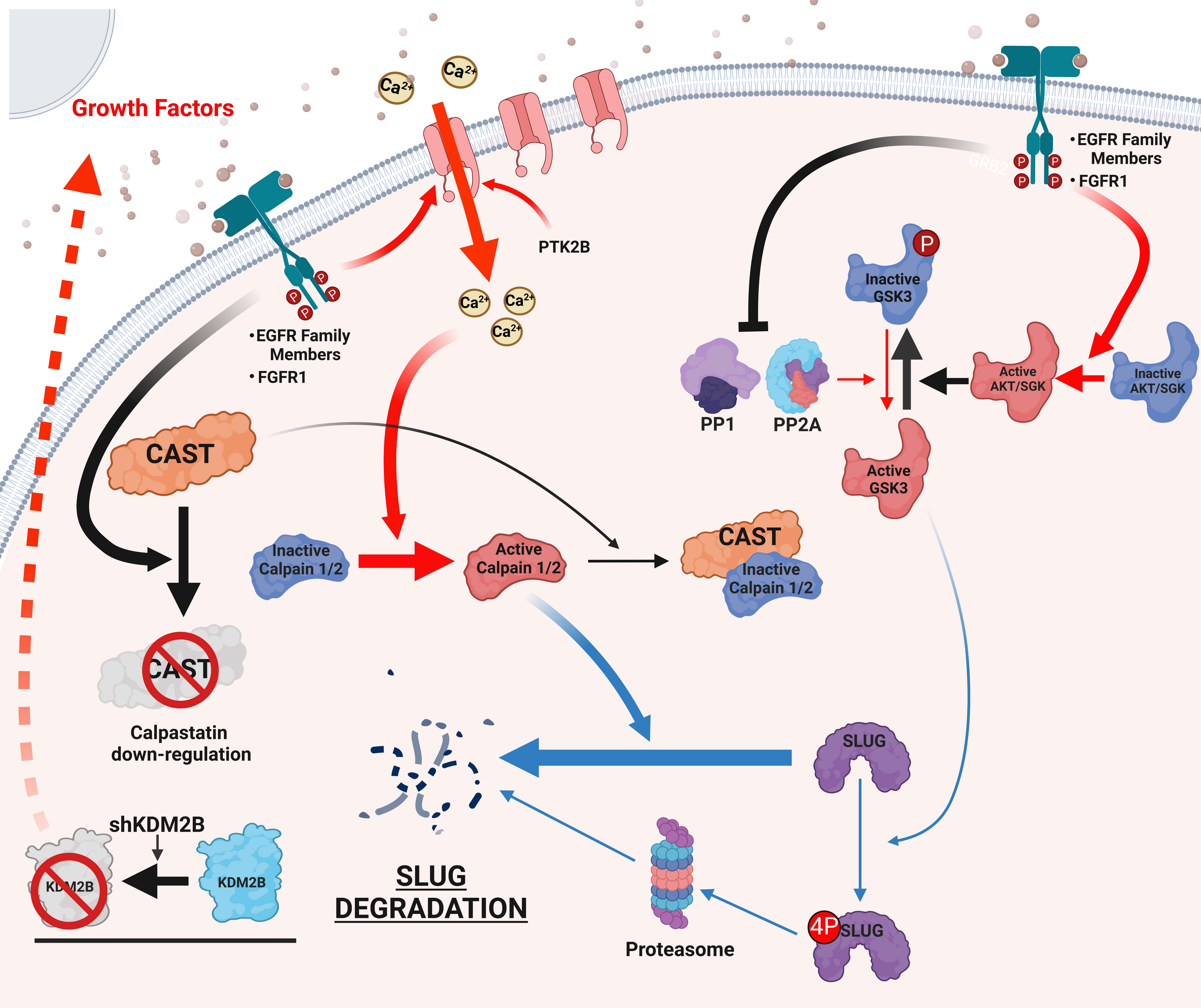

Growth Factors
Ca2+
PTK2B
•EGFR Family Members
•FGFR1
GRB2
Inactive GSK3
P
PP1
PP2A
Active AKT/SGK
Inactive AKT/SGK
Active GSK3
CAST
Inactive Calpain 1/2
Active Calpain 1/2
CAST
Inactive Calpain 1/2
CAST
Calpastatin down-regulation
SLUG
4P SLUG
Proteasome
SLUG DEGRADATION
shKDM2B
KDM2B
KDM2B
